# Folate deficiency disrupts key metabolic transitions within the developing neural ectoderm

**DOI:** 10.64898/2026.09.18.752622

**Authors:** Nicolas Dias, Daniel P. Lewinsohn, William N. Colgan, Minming Wang, Yusuke Kijima, JoAnne Villagrana, Tien-Chi Jason Hou, Gokul Gowri, Adelina Chau, Tuğçe Aktaş, Kaelyn Sumigray, Jonathan S. Weissman, Luke W. Koblan, Allon Wagner, Zachary D Smith

**Author notes:** Equal Contribution (Nicolas Dias, Daniel P. Lewinsohn). Equal Contribution (Luke W. Koblan, Allon Wagner, Zachary D Smith).

## Abstract

Despite long-standing epidemiological associations, the mechanism linking folate availability to gestational neural tube defects remains unclear, partly because measuring and interpreting metabolic activity in dynamic biological systems remains challenging. Here, we apply a deep-learning-based graph-guided variational autoencoder (MeRN; Metabolic Representation Network) to infer single-cell metabolic activity and states from scRNA-seq data of mouse embryogenesis. By analyzing folate-deficient embryogenesis from E7.0–E9.0, we identify a transient state within the nascent neural lineage that is acutely sensitive to folate availability, leading to an interconnected disruption between key bioenergetic pathways and *de novo* purine biosynthesis. Moreover, metabolically induced growth defects lead to permanent morphological disruptions along the dorsal-ventral axis, which we confirm by generating whole-embryo fate maps using a prime-editing-based lineage recorder (PEtracer). Collectively, our results establish a highly scalable framework for interpreting dynamic changes in embryonic metabolism and elucidating the mechanistic bases underlying environmentally linked congenital disorders.

**Research highlights:**

- Developing embryos adopt metabolic states that are distinct from cell lineage
- Metabolic states are organized temporally and spatially along key body axes
- Neural tube development depends on a rapid transition between metabolic states
- Activation of transient neural metabolic states is highly sensitive to disrupted folate absorption
- The floor plate is preferentially resistant to folate deprivation and distorts subsequent neural patterning

## Introduction

The well-established relationship between human neural tube development and maternal dietary folate highlights both the importance of metabolism during embryogenesis and the long-standing difficulty in defining how metabolic processes regulate tissue differentiation and morphogenesis^1–5^. Extensive clinical evidence links dysregulated folate metabolism to neural tube defects, prompting the fortification of staple foods with folic acid^2,6–9^. The consequent decline in the incidence of lethal or severe neural tube defects such as anencephaly and spina bifida is considered a landmark public health achievement of the 20th century, with reductions in incidence of ∼40-80% across adopting nations^2,9^. However, despite these striking epidemiological data, the mechanisms by which folate availability specifically regulates primary neurulation remain poorly understood, with hypotheses ranging from altered epigenome regulation to direct involvement in tissue biomechanics and folding^10–15^.

Folate, once metabolized to tetrahydrofolate (THF) and its derivatives, is the major carrier of one-carbon units in one-carbon metabolism (OCM)^16^. This metabolic pathway supports critical, diverse, and ubiquitous physiological processes including purine and thymidylate biosynthesis, S-adenosylmethionine recycling, and, as recently described, modulation of redox homeostasis through NADPH biosynthesis^17^. Given these essential roles in cellular physiology, distinguishing tissue-specific sensitivities from broader physiological requirements and system-wide stress responses to OCM disruption remains demanding, particularly given the difficulties in measuring metabolites in high-throughput single-cell assays^18^. Even with these measurements, interpreting differential metabolic activity is challenging given the scale and interconnectivity of biochemical reactions within individual cells. Understanding tissue-specific dynamics in OCM dependence would advance our understanding of how cellular metabolism influences growth, differentiation, and morphogenesis during early embryonic development. Further, these insights could clarify how metabolic dysregulation disrupts morphogenesis, identify diagnostic biomarkers of congenital disease, and reveal opportunities for novel therapeutic approaches^19,20^.

Despite broad interest in the metabolic regulation of embryogenesis, measuring and interpreting metabolic activity in developing systems faces substantial technical challenges. In part, the rapid changes in cellular composition and tissue diversity that characterize early developmental events are difficult to capture *in vitro,* even in advanced models such as embryoids or gastruloids^21–23^. Moreover, directly measuring metabolomes and metabolic fluxes in whole embryos remains infeasible in a comprehensive and scalable manner, particularly during highly dynamic events such as gastrulation or primary neurulation. Current single-cell metabolome approaches recover far fewer features than transcriptomic methods, with capture on the order of ∼100 measurements per cell across hundreds to thousands of cells, and even these limited data are difficult to connect to broader cell identity^24–27^. In contrast, single-cell RNA-sequencing (scRNA-seq), which captures transcriptional profiles of individual cells, has become standardized and highly scalable. Abundant, publicly available data have been generated to address diverse biological questions and assemble increasingly comprehensive atlases of animal development and disease^28–35^. As such, analytical methods that leverage existing transcriptomic data to infer single-cell-resolved metabolic activity provide a powerful framework for understanding the parameters that regulate complex developmental transitions and to explain the etiology of congenital abnormalities^36–40^.

Here, we apply the Metabolic Representation Net (MeRN) algorithm to study the metabolic regulation of embryonic growth and differentiation. Briefly, MeRN is a graph-guided variational autoencoder (VAE) that uses prior knowledge of metabolic topology to infer reaction activity from scRNA-seq data and embed this information into an interpretable, low-dimensional space (*Lewinsohn et al.*, accompanying manuscript). We use MeRN to construct a joint transcriptional and metabolic model of mouse gastrulation and organogenesis from a staged atlas of 72 whole mouse embryos, spanning E6.5–E9.0^41,42^. We identify cellular metabolic states throughout this developmental window, including a transient state highly enriched in the nascent neural lineage, supported by increased reliance on anaerobic energy metabolism and *de novo* purine biosynthesis. We show that metabolic states predict phenotypic abnormalities that emerge upon genetic disruption of the high-affinity folate receptor Folr1, including lineage-specific growth and morphological defects and distorted dorsal-ventral (DV) patterning. To assess the generalizability of this framework, we further apply MeRN to a cohort of 107 mutant embryos across 8 conditions associated with neural tube closure defects (NTDs) and use it to distinguish metabolic and non-metabolic genetic drivers of this common phenotype. Finally, we extend and validate these observations using PEtracer, providing, to our knowledge, the first use of an evolving barcode lineage tracing technology to quantify how an environmental perturbation alters progenitor pool sizes and dynamics during embryonic development^43^. Collectively, these results demonstrate that generative probabilistic models incorporating biological inductive biases can capture and interpret latent metabolic parameters of complex developmental processes, establishing a broadly applicable framework for connecting single-cell metabolic inference to the developmental origins of congenital disease.

## Results

### Characterization of metabolic activity in the developing mouse embryo with MeRN

We present MeRN, a graph-guided variational autoencoder (VAE) for inferring single-cell metabolic activity from scRNA-seq data, in an accompanying manuscript by Lewinsohn et al. Briefly, MeRN utilizes transcriptomic data and a predefined graph representation of the metabolic topology derived from the Kyoto Encyclopedia of Genes and Genomes (KEGG)^44^, where nodes represent biochemical reactions and edges represent shared metabolites. The model reconstructs single-cell gene expression profiles of the top 800 highly variable genes encoding metabolic enzymes (hereafter, “metabolic genes”) from a low-dimensional embedding (M), while simultaneously reconstructing non-metabolic (hereafter, “non-metabolic”) transcriptional variation from a separate embedding (**B**) (**Figure 1A**). To infer reaction usage, MeRN constrains the metabolic embedding via a specialized decoder that leverages the graph representation of metabolic reactions, first predicting each cell’s relative reaction activity (**R**) and then reconstructing the expression of related enzyme-encoding genes during training. Subsequently, MeRN outputs single-cell-level metabolic reaction activity and two low-dimensional embeddings that describe metabolic and non-metabolic cell-cell relationships, which can be leveraged for differential metabolic activity analyses, two-dimensional projection, cell clustering, and dynamic inference of metabolic transitions, among other applications (**Figure 1B**).

**Figure 1.**
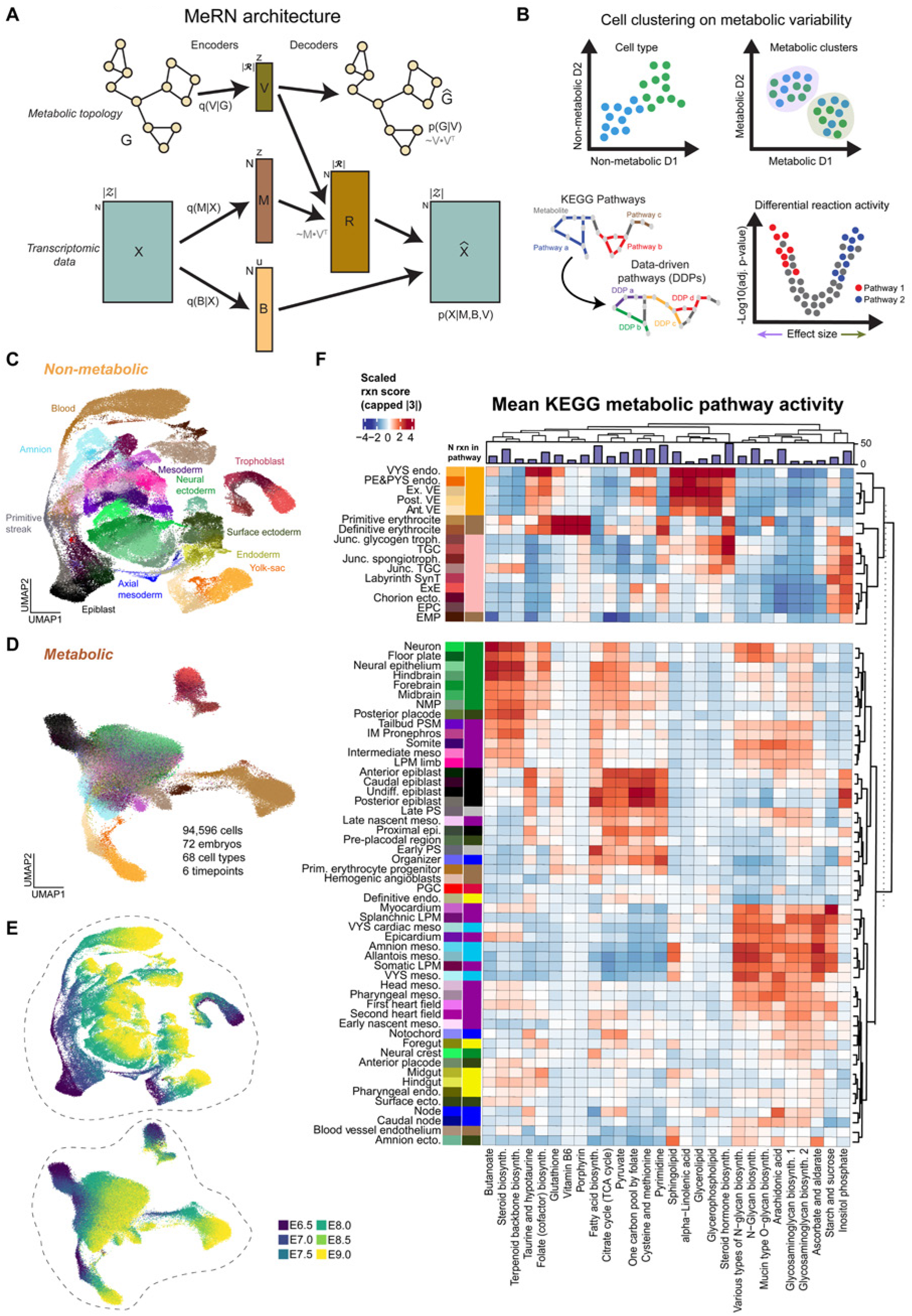
MeRN-based prediction of metabolic activity in whole embryos. **A.** MeRN model architecture. MeRN takes scRNA-seq data (X) and a KEGG-based graph representation of metabolic topology (G) as input. Neural network encoders (*q(V | G)*, *q(M | X)*, and *q(B | X)*) learn lower-dimensional embeddings for reactions (V) and cells, separated into metabolic (M) and non-metabolic (B) latent spaces. M is multiplied by *Vᵀ* to obtain the reaction activity matrix (R). R is decoded to reconstruct metabolic gene expression via a sparsified linear layer connecting reactions to transcripts encoding the enzymes for those reactions. Non-metabolic genes are decoded by a linear layer from B to *X̂*. Metabolic graph edges are decoded from pairwise dot products of the reaction embeddings in V. *p(G | V)* and *p(X | M, B, V)* denote the graph and scRNA-seq likelihood components. N denotes the number of input cells, z denotes the shared dimensionality of M and V, and u denotes the dimensionality of *B*. **B.** Downstream analysis utilizing MeRN output, including clustering using metabolic latent space, data-driven metabolic pathway calculation, and differential reaction and pathway activity. **C,D,E,** UMAP representation of the wild-type reference, calculated based on MeRN-derived non-metabolic (**C**) and metabolic (**D**) latent space embeddings, colored by cell types (**C**, **D**) and time points (**E**). **F.** Heatmap of the top 3 metabolic pathways per lineage, selected by Cohen’s d, highlights metabolic segregation into the gastrulation embryo and major nutrient support structures, including yolk sac (enriched for lipid biosynthesis pathways), placenta (enriched for hormone biosynthesis pathways), and primitive blood (vitamin B6 and porphyrin metabolism). The top bar plot highlights the number of reactions in each KEGG pathway. Annotation colors match those in (**C, D**). The score range is limited to an absolute value of 3 for visual clarity. Glycosaminoglycan biosynthesis sub-pathways are chondroitin/dermatan sulfate (**1**) and heparan sulfate/ heparin (**2**).

The developing embryo is a compelling use case for this method, as the complexity and dynamism of early embryogenesis severely hinder unbiased, scalable characterization of metabolic activity. Despite this complexity, developmental progress is relatively deterministic and robust to perturbation in standard conditions, providing a consistent reference outcome. To study embryogenesis, we first trained MeRN models on a recently published, densely sampled scRNA-seq staged atlas of wild-type mouse development, comprising 94,596 cells collected from 72 individual embryos and spanning 6 time points from E6.5 to E9.0 at 12-hour resolution^41^. This reference resolves 68 cell types from all major lineages, including the three embryonic germ layers and all extraembryonic tissues, using characteristic marker genes. We applied MeRN to compute latent metabolic and non-metabolic embeddings from these reference data (**Figures 1C, D, Figure S1A, B**). The non-metabolic embedding clearly delineates cell-type relationships and groups them into well-defined lineages. In contrast, the metabolic embedding separates pre- and post-gastrulation embryonic and extraembryonic lineages but conflates the three germ layers to varying degrees. This structure suggests that: (**1**) extraembryonic tissues have differentiable metabolic profiles, (**2**) embryonic cells proceed from, or tend to, a largely uniform metabolic state, and (**3**) the embryonic germ layers share core metabolic activities that distinguish them from the pluripotent epiblast, blood, and extraembryonic tissues. Notably, the latent geometry of both non-metabolic and metabolic embeddings preserved the temporal axis of embryonic development **(Figure 1E**), indicating that metabolic profiles in embryonic and extraembryonic tissues also depend strongly on developmental stage.

### MeRN identifies specialized metabolic features within nutritional support structures

To define each lineage’s characteristic metabolic activity, we calculated mean scaled KEGG pathway activation scores for each cell type by averaging their component reaction scores (**Figure 1F**). High-level k-means clustering (k = 2) clearly partitioned nutritional support systems such as the placenta, yolk sac, and blood from the epiblast and the three germ layers, which are comparatively more intermixed. We observed distinct reaction activity in the placenta, yolk sac, and blood, highlighting MeRN’s ability to capture interpretable, tissue-specific metabolic features from complex scRNA-seq data.

For example, the placenta and yolk sac are distinguished by increased activity in sphingolipid, glycerolipid, and glycerophospholipid metabolism (KEGG accessions **rn00600**, **rn00561**, and **rn00564**, respectively), in keeping with known functions that require frequent cell membrane remodeling and cell polarity for directional nutrient absorption, processing, and transfer to the embryo^45–49^. Placental cell types in the junctional zone also exhibit robust activity in steroid hormone biosynthesis (**rn00140**) (**Figure S1C, D**) and inositol phosphate metabolism (**rn00562**), reflecting their critical role in regulating the uterine environment^50,51^. Notably, our high-level clustering segregates primitive and definitive erythrocytes from less mature hemangioblasts and primitive erythrocyte progenitors, assigning functional red blood cells to the nutritional support structures. These cells exhibit increased activity in porphyrin (**rn00860**), vitamin B6 (**rn00750**), and glutathione (**rn00480**) metabolism pathways, which are associated with heme biosynthesis and hemoglobin assembly, and antioxidant defense, respectively^52–54^. We also observe high activity of pyrimidine metabolism (**rn00240**), particularly in reactions associated with ribonucleotide degradation via 5’-nucleotidases and dUTP diphosphatase (KEGG accession **R02100, R00511, R00512, R00515**) (**Figure S1E, F)**. This metabolic activity supports a model in which hypoxia-induced adaptation increases hemoglobin-oxygen affinity^55^, a feature of erythrocyte maturation previously described in embryonic red blood cells in chickens, and impaired in human disease^56,57^. Overall, these findings highlight MeRN’s ability to detect heterogeneous metabolic activity across and within cellular lineages, in keeping with known determinants of biological function.

### Identification of embryonic metabolic states

To better resolve metabolic state diversity within the embryo, we excluded extraembryonic endoderm and ectoderm, blood, and vascular cells from the data and trained MeRN models of gastrulation. To improve temporal resolution, we also ranked embryos from 1 to 72 by transcriptional similarity (hereafter, “embryo rank”; **Methods**). Leiden clustering of the resulting metabolic latent space revealed 6 general “metabolic states” (MS), each with its own temporal and cell lineage specificity (**Figure 2A, B, 2SA)**. For example, MS2 largely comprises early embryonic tissues such as the epiblast and primitive streak, while MS4 comprises most cardiac and extraembryonic mesoderm. In contrast, MS5, MS0, and MS1 consist of both neural and ectoderm cell types, as well as trunk mesoderm in varying proportions, while MS3 includes non-neural ectoderm, axial mesoderm, and endoderm (**Figure 2C**). Although transcriptional states are comparatively well defined and segregate into distinct lineages, cells with the same cell-type label were often assigned to multiple metabolic states, suggesting that metabolic features of cell identity may be more plastic. We also find that some metabolic states exhibit strong temporal dynamics. For instance, within the neural lineage, the Epiblast-enriched MS2 gradually declines in representation until E7.5, when MS5 supplants it, and peaks at E8.0 before MS0 rapidly replaces it at E8.5 (**Figure 2SB**). Dynamics within the neural lineage contrast those of MS3, MS4, and MS1, which emerge during gastrulation and are proportionally more stable once specified.

**Figure 2.**
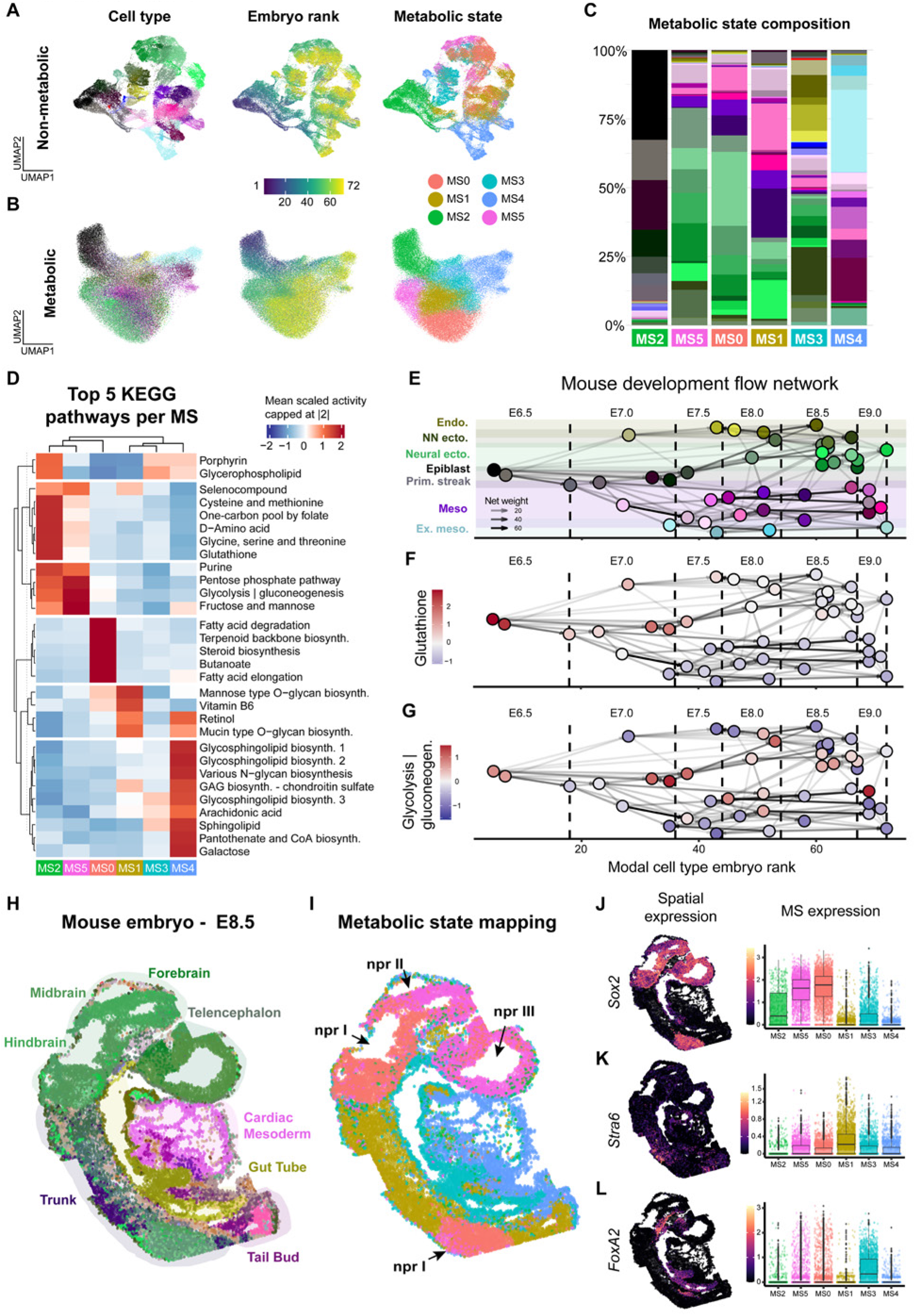
Metabolic state dynamics over mouse gastrulation. **A,B.** UMAP representation of embryonic lineages using MeRN-derived non-metabolic (**A**) and metabolic (**B**) latent embeddings. Cells are colored by cell type as in Figure 1, embryo rank (denoting relative developmental progress, **Methods**), and cluster (“metabolic state,” MS0-MS5). **C.** Bar plot showing average cell type composition per metabolic state, highlighting an enrichment for MS2 within the pluripotent epiblast and early gastrulation states, followed by stratification into neural/axial (MS5, MS0), mesodermal (MS1), endoderm/ectoderm (MS3), and cardiac/endothelial (MS4) enriched states**. D.** Heatmap of the top 5 metabolic pathways per metabolic state highlights the diversification of metabolic activity following gastrulation from the OCM-driven epiblast, including steady maintenance of these pathways through the transient Neural-enriched state MS5. Heat denotes mean pathway activity scaled across metabolic states. Rows are grouped by k-means clustering (*k* = 6). Heat is capped at an absolute value of 2 for visual clarity. **E,F,G.** Flow network graph layout of the control data^41^. Nodes represent cell types, and edges represent the maximum net transition probability between them. Nodes are colored by cell type labels (**E**) or scaled pathway activity for glutathione metabolism (**F**) and glycolysis and gluconeogenesis (**G**). The x-axis denotes the modal embryo rank per cell type. Guidelines represent embryo stages. Glutathione biosynthesis persists from the epiblast through to the early neural ectoderm before decaying in later states, while glycolysis and gluconeogenesis increase in relative activity within the same lineages. These dynamics are specific to these transitions, with activity decaying in other mesodermal and endodermal states. **H,I.** seqFISH data of a mouse embryo at E8.5, displayed in a sagittal section (replicate 2 from Harland et al.^75^). Cells are colored by cell type annotation (**H**) or predicted metabolic state (**I**). npr: approximate position of neuropores from neural tube closure sites I-III. Arrows show that these regions tend to represent boundaries between the transient neural state MS5 and the mature neural state MS0. **J,K,L**. Expression of representative genes within the spatial projection (left) and the average expression per metabolic state (right), with individual cells displayed. Neural tissues marked by *Sox2* (**J**), showing positions associated with the neural tube and heightened activity in MS5 and MS0. Trunk mesoderm marked by *Stra6* (**K**) is associated with positions posterior to the cervical region and MS1. Ventral position denoted by *FoxA2* (**L**) is associated with the endoderm, notochord, and ventral neural tube, and is enriched for MS3.

Comparative analysis of metabolic activity in each MS revealed defining characteristics, including the most active KEGG pathways and approximate relationships (**Figure 2D**). For example, the extraembryonic mesoderm-, vascular-, and cardiac-enriched MS4 is characterized by sphingolipid (**rn00600**), various forms of glycosphingolipid (**rn00601**, **rn00603**, **rn00604**), and arachidonic acid metabolism (**rn00590**). These pathways reflect active cellular signaling, particularly in the inflammatory response and in the regulation of vascular tone, and are likely connected to vasculogenesis in the developing endocardium^58–60^. The mesoderm-enriched MS4 and MS1 exhibit higher scores for retinol (**rn00830**) and glycan (**rn00515**, **rn00512**, **rn00532**) metabolism, indicating changes in cellular signaling associated with axial elongation and the acquisition of anterior-posterior (AP) positional identity^61–65^.

Compared with these starker transitions, the pluripotency and neural ectoderm-associated MS2 and MS5 states share substantial pathway activity in folate-dependent one-carbon metabolism (OCM), including the one-carbon pool by folate (**rn00670**), cysteine and methionine (**rn00270**), and glutathione (**rn00480**). Notably, these states also show several transitions related to energy metabolism, specifically the activation of glycolysis and gluconeogenesis (**rn00010**), the pentose phosphate pathway (PPP, **rn00030**), and purine metabolism (**rn00230**). We find that the pluripotency-enriched MS2 preferentially utilizes reactions linked to redox homeostasis, such as nicotinate and nicotinamide metabolism, whereas MS5 shows a relative increase in pathways responsive to hypoxia, such as sulfur metabolism^66^ (**Figure S2C)**. Given the temporal relationship between MS2 and MS5, these results likely reflect a collective shift from reliance on oxidative phosphorylation to a more glycolytic state between ∼E7.5 and E8.0, consistent with the role of hypoxia in neural tube formation and changes to energy metabolism during neural tube morphogenesis^67–72^. To better characterize this transition, we calculated a flow network based on our cell-type annotation, defining likely developmental transitions between cell types^41,73^ (**Figure 2E**). We further calculated the mean scaled activities of glutathione and glycolysis/gluconeogenesis for each cell type and projected these scores onto the network (**Figure 2F, G**). We find that predicted glutathione activity is higher before E7.5, and glycolysis increases at later time points, particularly in the neural ectoderm. Because glutathione metabolism responds to non-oxidative factors, we also analyzed the average expression patterns of 143 hallmark hypoxia genes^74^ (**MM3861**) over time and found they were also upregulated between E7.5 and E8.0 (**Figure S2D**, **Table 1**). Collectively, these data indicate a transient shift towards a glycolytic state during early neural lineage specification, as captured in our MS5 annotation.

### Metabolic states conform with positional identity and neural tube morphodynamics

Because spatial position is a central determinant of cell fate during embryogenesis, we next sought to resolve the metabolic states identified by MeRN within their anatomical context. Using seqFISH data reported by Harland et al^75^., we predict metabolic states for cells within sagittal sections of mouse embryos at E8.5 according to our MeRN model (**Figure 2H, I; Methods).** Overall, metabolic states are strongly associated with axial position and broadly consistent with developmental lineages, including enrichment of MS4 within the cardiac mesoderm, ventral enrichment of MS3 at the expected position of the gut tube, and medial-caudal enrichment of MS1, posterior to the cervical region and extending down to the tail bud. The neural lineage-associated states, MS5 and MS0, are primarily positioned anterior to the cervical-hindbrain boundary. Importantly, these trends held across biological replicates, and the relationship between metabolic states and cell types is consistent with our previous findings from our orthogonally generated data (**Figures S2E-I**).

Neural-associated metabolic states occupy distinct spatial domains in the developing embryo, with the boundary between MS5 and MS0 aligning with known morphological transitions during neural tube development (**Figure 2J**). Specifically, cells in the anterior-dorsal regions of the embryo, proximal to neuropores (npr) II and III, near the forebrain-midbrain boundary, and in the rostral telencephalon consistently map to MS5. In contrast, cells near the anterior and posterior closure sites for npr I preferentially map to MS0, supporting a likely MS5-to-MS0 transition during neural tube closure. This dynamic is bounded by the MS1-associated trunk mesoderm, which spans the edges of neural tube closure I sites, and exhibits heightened retinoic acid metabolism, as inferred from our MeRN results and consistent with a medial position roughly demarcated by *Stra6* expression. (**Figure 2K**). While our model does not define characteristic metabolic activity for MS3 outside of cell membrane-associated pathways, its spatial positioning indicates a strong ventral identity, particularly associated with the developing gut tube and *FoxA2* expression (**Figure 2L**). This association is consistent with our data and, combined with its proximity to the neural tube, suggests that MS3 may also be related to ventralizing morphogenetic signaling^76^.

### Defective folate transport impacts neural and axial growth

Our analysis of MeRN-inferred metabolic activity revealed a key neural-specific metabolic state transition, from MS2 to MS5, associated with the earliest stages of neural fate induction and defined by a co-occurring shift to hypoxia-like metabolism and high OCM-associated pathway utilization. However, how folate deprivation may act at the interface of these two metabolic states to disrupt neural tube closure remained unclear. To evaluate the effects of folate deficiency in early embryogenesis, we genetically disrupted the principal dietary folate receptor (*Folr1*) in mouse embryos using a zygotic CRISPR/Cas9 knockout (Crispant) approach^31,34,77–81^ (**Figure S3A**, **Methods**). This genetic model (hereafter, Δ*Folr1*) limits variability inherent to maternal diet and drug-based interventions, providing a highly controlled setting to study folate metabolic disruption, as mice lacking functional *Folr1* display hallmark signs of high-penetrance folate deficiency^82–85^. The morphological phenotype at E9.0 for Δ*Folr1* embryos includes reduced embryonic size, recessed surface ectoderm surrounding the head fold, disrupted neural tube closure and cardiac morphology, and loss of somite structural definition (**Figure 3A, B**). Boundaries between somites are less well-defined but are generally preserved, indicating that some AP patterning is retained despite the lack of axial rotation in the caudal embryo. However, Δ*Folr1* somites lack a characteristic actin-rich medial domain, again suggesting a downstream DV patterning defect^86^.

**Figure 3.**
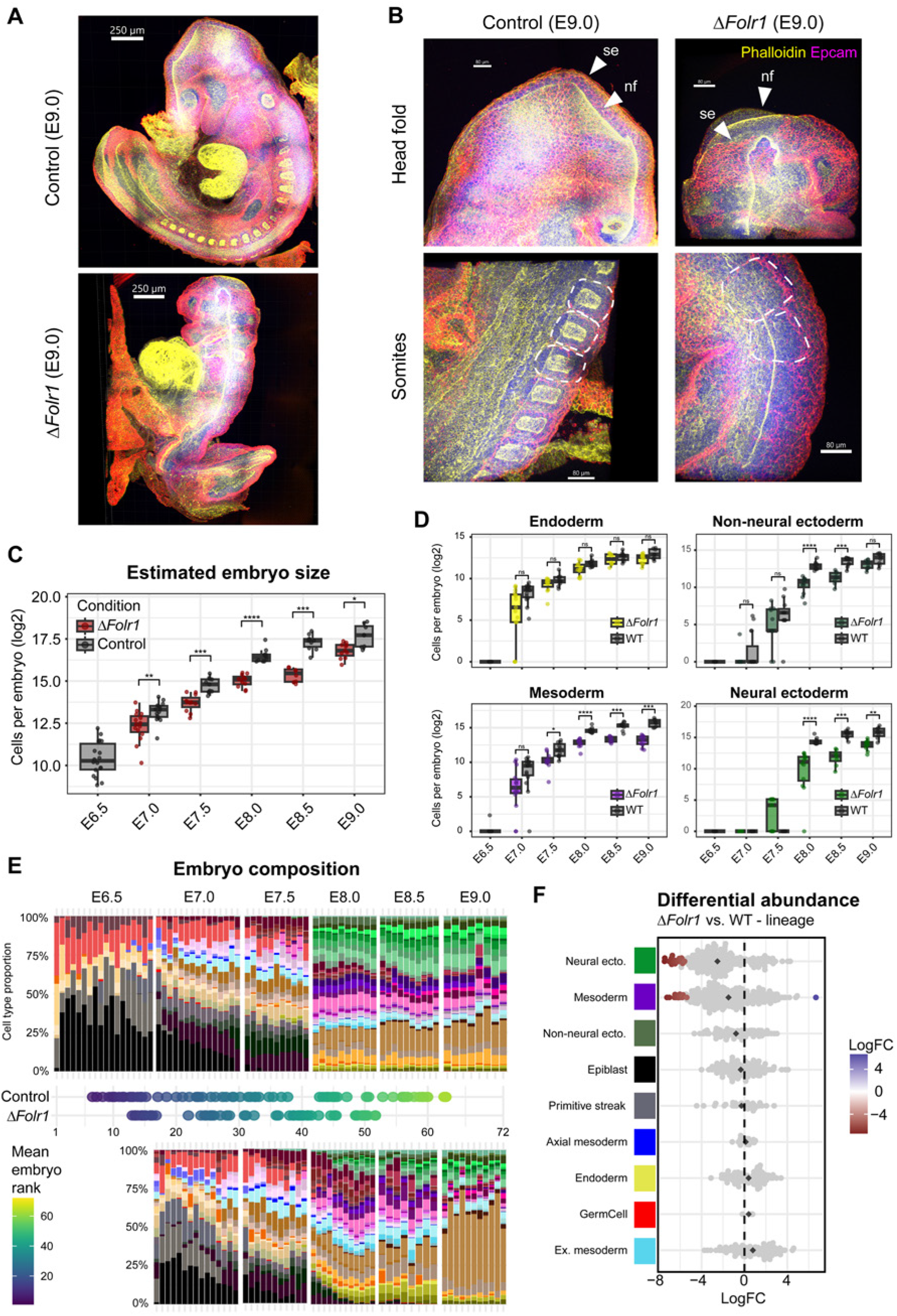
*Folr1* mutant embryos exhibit lineage-specific growth defects. **A.** Representative whole-mount images of E9.0 embryo morphology in sagittal view. Control (top) and Δ*Folr1* (bottom) conditions are displayed. F-actin stained by phalloidin (yellow) and epithelial cells stained by Epcam (magenta). Scale bars: 250 μm. **b.** Zoom-in view of neural tube morphology, including the head fold (top) and somites (bottom) in the wild-type control (left) and Δ*Folr1* (right) conditions. Δ*Folr1* embryos display characteristic features of NTDs, including exposed and open neural epithelium, as well as recessed surface ectoderm. Although somites form, they lack characteristic definition, including strong apical F-actin enrichment. Dashed lines demarcate approximate somite boundaries. se: surface ectoderm, nf: neural fold. Scale bars: 80 μm. **C,D.** Boxplot of log2-transformed cell count estimates per embryo as inferred from scRNA-seq of 57 Δ*Folr1* embryos collected from E7.0 to E9.0, grouped per stage and split by condition. Samples represent whole embryos (**C**) or individual lineages (**D**), including endoderm, non-neural ectoderm, mesoderm, and neural ectoderm. Growth disparities in Δ*Folr1* embryos increase over time and are preferential to the neural ectoderm and mesoderm compared to mesodermal cell types. Significance was calculated using a multiple-hypothesis-corrected (Bonferroni) one-sided Mann-Whitney U test (hypothesis: Δ*Folr1* embryos are smaller than control). **E.** Cell type composition for individual control (top) and Δ*Folr1* (bottom) replicate embryos, grouped by collection time point. Mean embryo rank was calculated by averaging predicted ranks per cell based on the wild-type reference (see Methods). **F.** Differential abundance results from Milo, using the non-metabolic latent space and grouped by lineage, confirm that all major cell types are produced over gastrulation, with significant depletion of select mesodermal and neural ectodermal cell types. Points represent neighborhoods; color denotes significance (spatial FDR < 0.05). Black diamonds represent group means.

For comparative analyses with our scRNA-seq wild-type reference, we recovered and profiled 57 embryos collected between E7.0 and E9.0 with 12-hour resolution, capturing 57,034 viable transcriptomes (**Figure S3B**). By combining cell counting with our demultiplexing strategy, we find that Δ*Folr1* embryos show a progressive growth defect, ranging from an average 0.64-fold decrease in cell number at E7.0 to 0.58-fold at E9.0, with the lowest difference at E8.5 (0.29-fold) (**Figure 3C, Methods**). All 68 wild-type cell types are represented, albeit in different proportions, suggesting that the differentiation events associated with gastrulation nonetheless occur in the absence of efficient dietary folate capture (**Figure 3D, E, S3C, D**). Extending the estimated cell counts per embryo analysis to individual lineages revealed specific patterns of lineage-specific disruption by folate availability, particularly in the neural ectoderm and axial mesoderm (**Figure 3D**). In contrast, we observed no significant depletion in the embryonic endoderm, consistent with the previously characterized and reported neural tube-specific phenotype^82^.

To better understand the temporal dynamics of embryo composition, we ranked ΔFolr1 samples by transcriptional similarity and analyzed cell-type composition in a near-continuous fashion. This analysis revealed a general developmental delay in cell-type proportions, particularly evident during the transition from E7.5 to E8.0, a period associated with dramatic embryonic growth and increasing embryonic complexity (**Figure 3E**). Mapping our ΔFolr1 cells to our non-metabolic latent space and measuring differential abundance with Milo^87^, an overlap-aware KNN-based approach, revealed that few neighborhoods showed significant compositional differences, primarily characterized by depletion in the neural ectoderm and mesoderm lineages (**Figure 3F, S3E-G, Methods**). More specifically, neighborhoods with significant depletion of Δ*Folr1* cells (n = 29 of 873 neighborhoods) include somites, neural-mesodermal progenitors (NMPs), and fore-, mid-, and hindbrain (**Figure S3H**). Notably, a single neighborhood, designated as splanchnic lateral plate mesoderm (LPM), shows a significant enrichment for Δ*Folr1* cells. This tissue represents the ventral-most portion of the outer layer of the neural tube-adjacent mesoderm, and its dorsal counterpart, the somatic LPM, is generally depleted. These results agree with the morphological phenotype and further suggest a downstream DV patterning defect. Importantly, while Δ*Folr1* embryos appear to exhibit differential, cell-type-specific growth defects, they do not appear to affect the forward differentiation of depleted states.

### Metabolic state membership is acutely connected to folate sensitivity

While the observed cell-type depletion pattern broadly matched the morphology of Δ*Folr1* mutants, the lack of cell-type specificity or complete ablation suggests that disruption of folate metabolism does not uniformly affect cells within a lineage. We then hypothesized that metabolic state preference, rather than lineage, could be a more robust unifying factor in cell depletion. Indeed, the predicted metabolic embeddings and metabolic state assignments for cells in Δ*Folr1* embryos immediately revealed markedly different occupancy in the metabolic low-dimensional projection (**Figure 4A**; **see Methods**). In particular, the region populated by E8.0 cells belonging to MS5 appeared substantially depleted, which Milo analyses confirmed as a robust and highly significant effect (**Figure 4B, C, S4A**). Of the 848 neighborhoods assayed, 339 show significant differential abundance, with most (67.8%) representing Δ*Folr1*-depleted regions. Following neighborhood annotation, we found that while MS2, MS3, and MS4 show little change in abundance, 98.0% of neighborhoods dominated by MS5 cells were significantly depleted in Δ*Folr1* embryos (average log-fold change = −5.46), a stronger and more coherent effect than observed for any individual cell type or lineage (**Figure 4C**, **S4B**). We observed a significant and robust negative correlation between MS5 membership and relative cell-type abundance at E8.0, the period of maximal MS5 representation, confirming preferential depletion of cells in this state (R = -0.65, p = 3.7e-4, **Figure 4D**). Conversely, aggregate membership in MS3 and MS4 was significantly associated with enrichment and predominantly highlights cell states with strong ventral identity, such as the caudal node, the embryonic gut, and the splanchnic lateral plate mesoderm (R = 0.75, p = 1.2e-5, **Figure 4E**). As such, metabolic state membership appears prognostic for sensitivity to maternal dietary cofactors, and MS5 shows both high OCM-related metabolic activity (**Figure 2D**) and susceptibility to folate metabolism disruption (**Figure 4C**).

**Figure 4.**
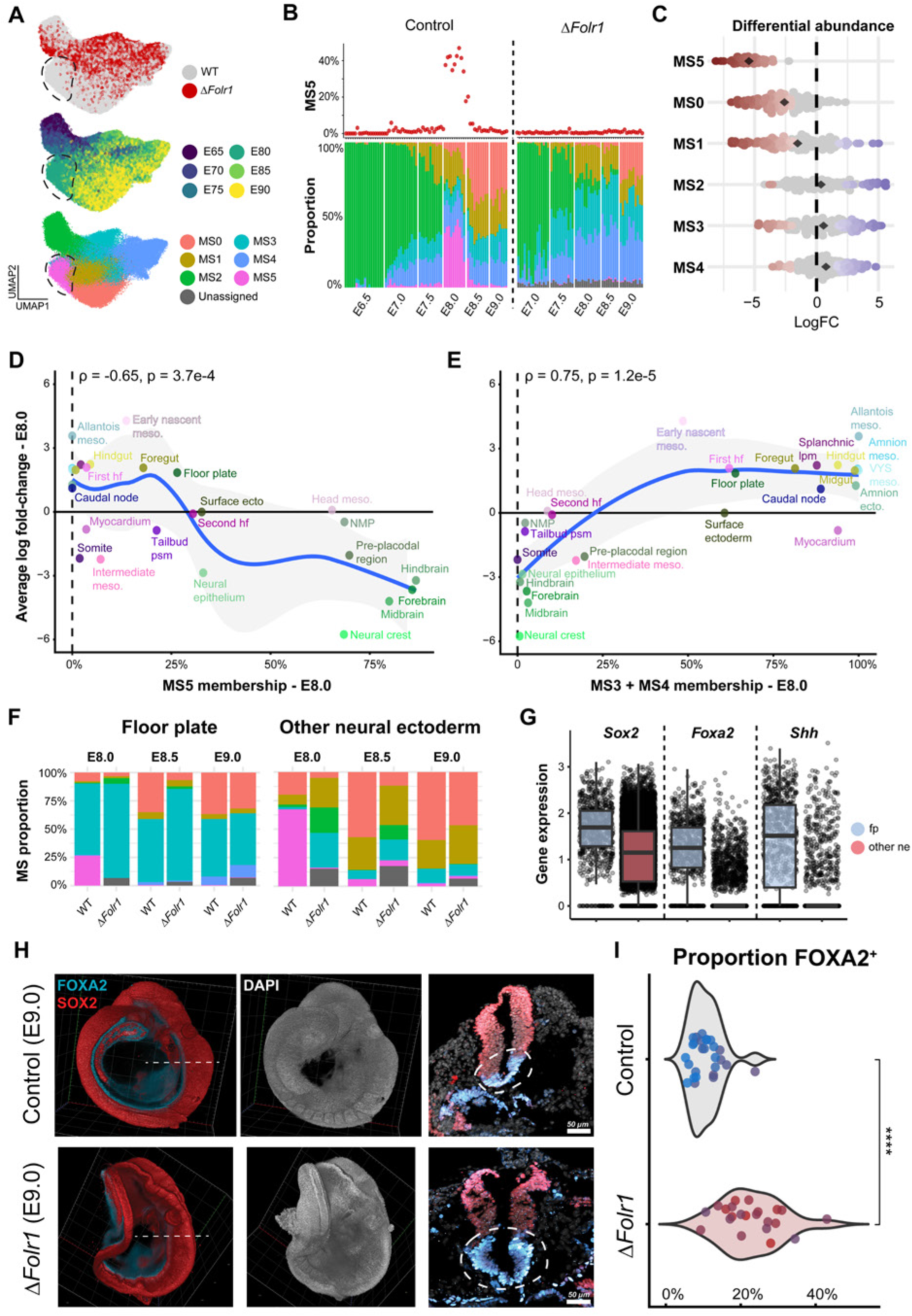
Disrupted folate transport blocks OCM-dominant metabolic state transitions. **A.** Joint UMAP representation of control and Δ*Folr1* data colored by condition, stage, and metabolic state. The dashed line represents the area occupied by E8.0 and MS5, and lacking Δ*Folr1* cells. **B.** Metabolic state composition throughout development. Bars (bottom) represent metabolic state proportions per embryo, and points (top) represent the proportion of MS5 cells. Unlike the non-metabolic transcriptional embeddings, the MS5 state is never substantially induced over Δ*Folr1* gastrulation. **C.** Differential abundance results from Milo using the metabolic latent space confirm extreme depletion of MS5 and secondary depletion of MS0. Points represent neighborhoods; color denotes significance (spatial FDR < 0.05). Black diamonds represent group means. **D,E.** Relationship between metabolic state membership in control embryos and cell type differential abundance in E8.0 Δ*Folr1* embryos (average Log2-fold change). MS5 (**D**) shows a significant negative correlation, indicating a clear relationship between MS5-encoded reactions and folate-dependent growth, while MS3 and MS4 (**E**) are comparatively resistant. Lines generated with Loess smoothing regression; correlation coefficient (R) and significance calculated with Spearman correlation. **F.** Average metabolic state composition of the floor plate compared to all other neural ectoderm states, split by timepoint, and grouped by condition. Bar slices are colored by metabolic state, as in (**A)**. **G.** Expression of floor plate marker genes between floor plate (blue) and other neural ectoderm cells (red). **H.** Representative whole-mount embryos (left) and frontal sections of the neural tube (right) co-stained with SOX2 (neural) and FOXA2 (ventral) antibodies to determine the relative dorsal-ventral identity of the neural tube along the anterior-posterior axis; co-expression of SOX2 and FOXA2 marks the floor plate. **I.** Quantification of the percent FOXA2^+^ floor plate within the SOX2^+^ neural tube for three representative embryos. Individual colors represent measurements from replicate embryos, with sampling at multiple points along the anterior-posterior axis. Significance was calculated with Welch’s t-test (p-value = 3.19e-07).

### Differential folate sensitivity skews dorsal-ventral patterning

While metabolic states predicted differential growth, MeRN also points to an exception in neural lineage depletion. Specifically, the floor plate exhibits both low MS5 membership and overall enrichment in Δ*Folr1* embryos, in contrast to all other neural ectoderm cells captured. A closer investigation revealed that the floor plate is substantially enriched for MS3, which is otherwise associated with the endoderm and notochord, two cell states that are also comparatively resistant to defective folate transport (**Figure 4F**). This pattern was consistent across our captured stages and was further enhanced in ΔFolr1 embryos, supporting the link between metabolic state and differential abundance.

Morphologically, the floor plate is the ventral-most portion of the neural tube and functions as a ventralizing organizer through Shh signaling, superseding the caudal node and notochord in this role^88–91^ (**Figure 4G**). Physiologically, the floor plate also differs from the rapidly expanding neural tube by its relative quiescence, where tissue expansion largely occurs through morphogenetic patterning rather than cellular proliferation^92,93^. Consistent with this, cell cycle predictions using the computational method tricycle^94^ confirmed that the slower-cycling floor plate and post-mitotic neurons are generally unaffected in Δ*Folr1* embryos, whereas proliferative neural ectoderm, such as Neuromesodermal Progenitor (NMP) cells and neural epithelium, preferentially stall in S or G0 phase, an expected result of nucleotide insufficiency (**Figure S4C-E**). Overall, these results suggest that the floor plate’s metabolic characteristics explain its resistance to folate deprivation, despite its neural lineage, and this resistance could naturally skew the DV axis by promoting excessive ventralization through Shh signaling. To validate these findings *in situ*, we investigated neural tube patterning in developing embryos at E9.0 using light-sheet microscopy to capture the DV axis of the neural tube from anterior to posterior (**Figure 4H, S4F, G1**). We observe consistent expansion of FoxA2^+^ neural ectoderm throughout the embryo, consistent with the abnormal floor plate expansion projected by our computational analyses, and with known biology of floor plate induction^95^ (**Figure 4I**).

### Critical breakdown between glycolysis and purine metabolism

Our results indicate that loss of dietary folate uptake significantly depletes a transitional metabolic state associated with neural development at the onset of primary neurulation (MS5). However, the metabolic activity underlying this dynamic remained unclear and difficult to describe using pre-existing pathway annotations. While canonical metabolic pathways are generally descriptive, they may overlook context-specific metabolic configurations and obfuscate functional links between reactions across different pathways. To address this, we applied a metabolic data-driven pathway (DDP) approach, introduced in the accompanying manuscript by Lewinsohn et al. (**Figure 5A**). Briefly, this method is agnostic to prior annotations and instead leverages the predefined topology of the KEGG metabolic reaction graph and MeRN-predicted reaction activity to identify functionally connected pathways based on direct connectivity and covariation within the data (**see Methods**). To focus our analysis on differences in folate sensitivity, we calculated DDPs using MS2 cells exclusively, reasoning that its role as a “progenitor” state would reflect the dysregulated metabolism that leads to NTDs upon attempted entry into MS5 (**Figure 4A, B**). This approach yielded 129 DDPs, with an average of 8 positively correlated and interconnected reactions per pathway (**Table 2**).

**Figure 5.**
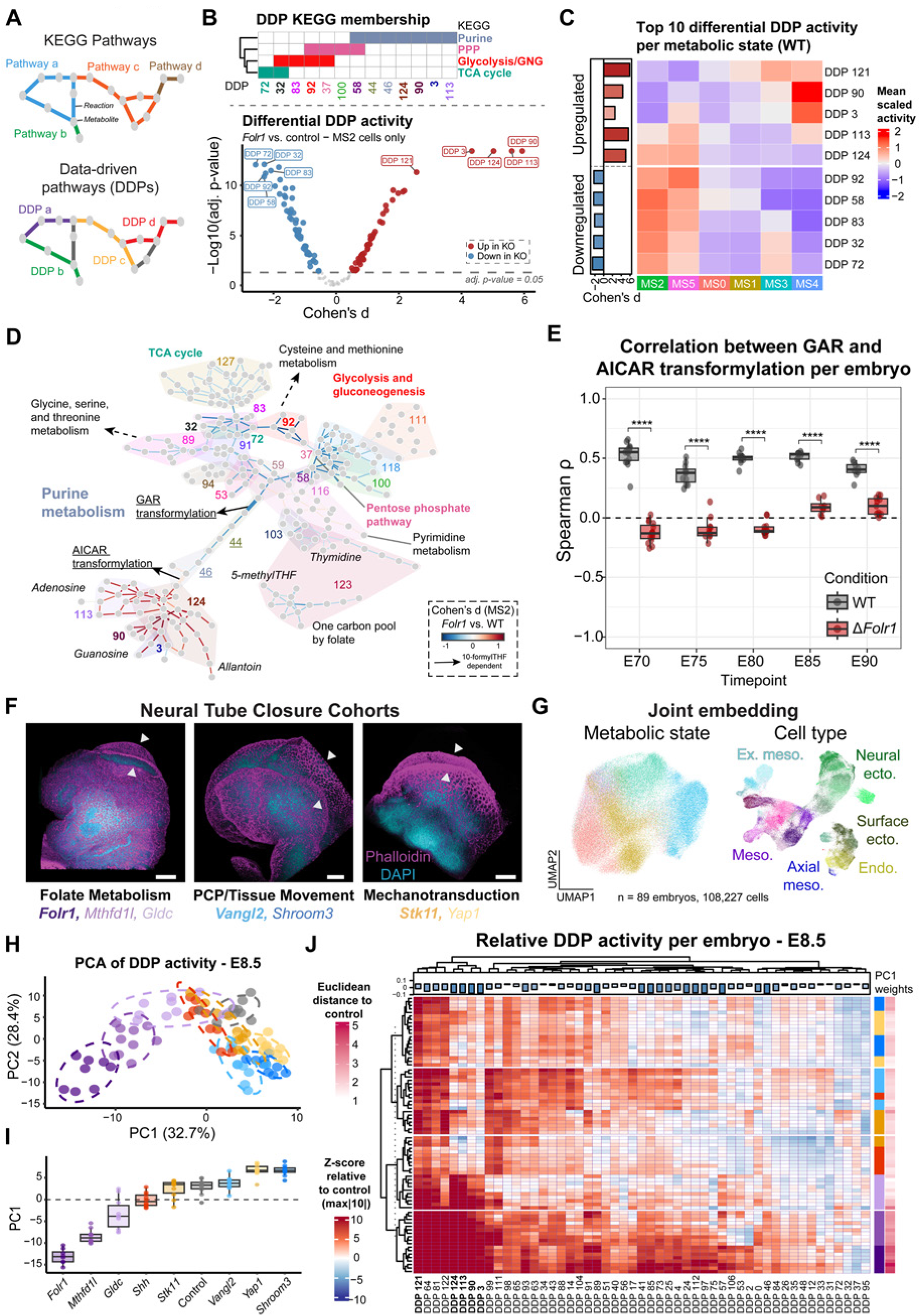
Metabolic NTDs reflect altered flux between glycolysis/gluconeogenesis and purine biosynthesis. **A.** Schematic of the Data-Driven Pathway (DDP) based approach for grouping reactions into biochemically defined pathways (see the accompanying manuscript, Lewinsohn *et al.*). Colors represent KEGG pathway (top) or DDP groupings (bottom)**. B.** Differential DDP activity between control and Δ*Folr1* MS2 cells, pseudo-bulked per replicate embryo. Color denotes significance (α < 0.05) and either upregulation (red) or downregulation (blue) in the experimental condition. The top 5 pathways per direction are highlighted. Displayed p-values were calculated with a two-sided Mann-Whitney U test and adjusted with the Benjamini-Hochberg correction. The annotation heatmap denotes membership in selected KEGG pathways, highlighting connectivity between energy metabolism and purine biosynthesis among dysregulated DDPs. **C.** Heatmap of the top 10 upregulated and downregulated DDPs displayed in (**B**). Heat denotes the mean activity per metabolic state, scaled across groups. Annotation denotes Cohen’s d displayed in (**B**). **D.** Metabolic network representation of reactions involved in energy and folate-related metabolic pathways, where nodes represent metabolites and edges represent reactions. The top significantly dysregulated DDPs are highlighted in Bold. Edge colors denote Cohen’s d for each reaction between all Δ*Folr1* and control cells. Reactions grouped into key KEGG pathways are annotated. Arrows denote 10-formyltetrahydrofolate-dependent reactions in purine metabolism within two connected DDPs: **DDP 44 (downregulated)**, containing an upstream reaction that gates purine metabolism from the pentose phosphate pathway by synthesizing FGAR from GAR (produced by phosphoribosyl pyrophosphate or ribose-5-phosphate from the PPP and ADP-ribose degradation, respectively); **DDP 46 (upregulated),** containing a downstream reaction that utilizes AICAR to enable forward progression into inosine monophosphate and *de novo* purine biosynthesis. **E.** Boxplot of Spearman correlation between activity in 10-formyl-tetrahydrofolate-dependent reactions in purine metabolism highlighted in (**D**), in wild-type (gray) and Δ*Folr1* (red). Each dot represents an embryo. Significance was calculated with a two-sided Mann-Whitney U test and adjusted with the Benjamini-Hochberg correction (****: *p_adaa_* < 3.0*e* − 5). **F.** Representative 3D rendering of NTD mutant cohorts, including those associated with folate transport and metabolism (*Folr1*, *Mthfd1l*, *Gldc*), planar cell polarity (PCP) and tissue movement (*Vangl2*, *Shroom3*), and mechanotransduction (*Stk11*, *Yap1*). We also included Δ*Shh* mutants to model loss of the ventralizing signal. Highlighted labels indicate the condition shown. White arrows highlight the neural folds and unclosed neural tube. Scale bars: 100 µm. **G.** UMAP representation of merged mutant and wild-type scRNA-seq data at E8.5, as calculated using the metabolic latent space with metabolic state annotation (left) and the non-metabolic latent space with cell type annotation, as shown in **Figure S5D**. **H, I.** Principal component boxplot (**I**) and biplot (**H**) showing the principal component position of individual embryo replicates calculated from MeRN-predicted DDP activity. Individual points represent embryos, colored by the target gene**. J.** Heatmap of DDP activity relative to the control data shows that less penetrant folate metabolism genes affect the same pathways as predicted for Δ*Folr1*, though often less completely and to a lesser degree. Rows denote individual embryos, grouped according to k-means clustering for clarity (*k* = 4). Row annotations (right) indicate the mutant condition and the Euclidean distance from the control, based on average DDP activity. Column annotation denotes the weight of each pathway based on the PCA calculated in **H**.

To evaluate differential activity between conditions, we calculated Cohen’s d for each DDP between control and Δ*Folr1* MS2 cells, averaging activity scores per embryo to mitigate pseudo-replication (**Figure 5B**). Although we found no evidence for MS2 depletion in Δ*Folr1* embryos, we identified several DDPs already differentially active in this state, suggesting a broad metabolic disruption that precedes primary neurulation and may link to interference in MS5 induction. Notably, most of these DDPs were preferentially active in MS2 and MS5, further supporting the functional link between these states and the role of this connection in folate sensitivity (**Figure 5C**). Visualization of the underlying connections between metabolic reactions revealed that the four most upregulated pathways in Δ*Folr1* are immediately downstream of a 10-formyltetrahydrofolate-dependent commitment step for *de novo* purine biosynthesis (DDP 46, **R04560, Figure 5D**). Conversely, downregulated DDPs were interconnected upstream of a second folate-dependent reaction, capturing a path that included glycolysis and the TCA cycle, as well as the PPP before converging on purine metabolism. This pattern created an interesting dynamic in which Δ*Folr1* embryos simultaneously downregulate energy metabolism and the PPP while upregulating folate-independent steps in downstream *de novo* purine biosynthesis, potentially as a compensatory increase in nucleotide production efficiency in a folate-scarce environment. Notably, our results do not suggest similar dysregulation in other aspects of OCM – such as glycine, serine, threonine, cysteine, or methionine metabolism – further supporting the functional link between glycolysis and purine biosynthesis as a determinant of successful neural growth and differentiation (**Figure S5A, B).** Combined with the wild-type activity pattern and the state-specific depletion of MS5, these data suggest that purine metabolism critically regulates entry into MS5, particularly through its folate-dependent reactions.

To explore this dynamic further, we investigated the activity of GAR and AICAR transformylation reactions (**R04325** and **R04560**, respectively), intermediate steps in *de novo* purine biosynthesis that require a 10-formyltetrahydrofolate equivalent and connect glycolysis and the PPP to purine metabolism. To do so, we trained MeRN models using the embryonic subset of the Δ*Folr1* data and compared the correlation between these two reactions’ activities across cells within each embryo. We found a consistent, significant loss of correlation at all time points, indicating a sustained breakdown in the chain of reactions that connect energy metabolism to *de novo* purine biosynthesis (**Figure 5E**). Importantly, this pattern repeated across metabolic states, indicating that embryo composition does not substantially drive this effect (**Figure S5C**). Instead, this decoupling between major energetic and biosynthetic pathways appears to reflect a critical breakdown in the metabolic requirements for rapid expansion of the neural epithelium.

### MeRN distinguishes metabolic and non-metabolic drivers of NTDs

While folate availability is strongly linked to early neural development, neural tube abnormalities do not respond uniformly to folate supplementation, nor is their pathogenesis solely linked to one-carbon metabolism^96–99^. We therefore reasoned that MeRN may deconvolute metabolic and non-metabolic sources of neural tube closure defects (NTDs). To assess converging and diverging characteristics of NTD-associated mutations, we generated additional scRNA-seq data for embryos at E8.5 with a diverse array of perturbations, including additional factors associated with one-carbon metabolism^100,101^ (*Gldc*, 10 embryos, *n* = 10,419 cells; *Mthfd1l*, 10 embryos, *n*= 17,981 cells), cell morphology and membrane polarity regulators^102–105^ (*Vangl2*, 10 embryos, *n*= 28,363 cells; *Shroom3*, 10 embryos, *n* = 27,423 cells), and mechanotransduction^106,107^ (*Yap1*, 10 embryos, *n* = 22,995 cells; *Stk11*, 10 embryos, *n* = 25,887 cells). These mutant embryos show abnormal neural tube morphology, with varying penetrance and effects on other tissues. Additionally, as a contrast to the observed ventralizing effect of *Folr1* disruption, we generated samples lacking the floor plate by perturbing *Shh*, a critical morphogenetic factor associated with ventral development^108^ (10 embryos, *n* = 18,192 cells) (**Figure 5F, S5G)**.

For comparative analysis, we applied our embryonic MeRN models to the additional mutant data, predicting metabolic and non-metabolic latent space embeddings as well as reaction activity and metabolic state assignments (**Figure S5D-F, Methods**). To summarize metabolic activity variance across conditions, we performed a principal component analysis of DDP activity per embryo, restricting it to E8.5 as the common stage (**Figure 5G-I**). The largest source of variance, represented by PC1 (32.7%), clearly separated Δ*Folr1* samples from other mutant embryos and functionally aligned our other metabolic perturbations by phenotypic severity (**Figure 5I**). To better understand the metabolic activity driving this partitioning, we examined the relative activity of DDPs with negative PC1 weights compared to controls (**Figure 5J**). This analysis revealed that purine metabolism-associated DDPs (124, 113, 90, 3) are similarly dysregulated across all folate-associated mutants but not in non-metabolic samples, supporting our conclusions regarding the link between folate metabolism and NTD etiology.

We also identified pathways upregulated across all assayed mutant cohorts, defined by DDPs 121, 64, 81, and 122 (**Figure 5J**). Inspection of their connectivity revealed a shared functional relationship to arachidonic acid metabolism, a polyunsaturated acid associated with the inflammatory response and recently proposed as a serum biomarker for neural tube defects^109^ (**Figure S5H**, **Table 2**). Our results corroborate this clinical observation and provide supporting evidence for arachidonic acid as a general marker for early identification of congenital NTDs.

### Fate maps of *Folr1* mutant embryogenesis

Our inference of metabolic activity in mouse gastrulation predicts that *de novo* purine metabolism is strained during the transition from epiblast to early neural ectoderm, increasing sensitivity to folate availability in these lineages. To confirm that the resulting phenotype reflects disrupted growth of these cell types, rather than biased cell fate specification, we used the PEtracer lineage recorder^43^ to reconstruct developmental cell-division trees under folate-depleted conditions. We injected Cre-activated mouse embryonic stem cells containing the PEtracer system into Δ*Folr1* morulae and recovered biological triplicate embryos at E9.5 to compare to previously generated E7.5-E10.0 reference data^43^ (**Figure 6A-D, SA-C, E**). ). All Δ*Folr1*-incorporated PEtracer embryos displayed near-identical morphological features and similar lineage composition to the zygotic Δ*Folr1* embryos described above, despite being composed mostly of *Folr1^+/+^* donor cells, suggesting that defective folate capture from Δ*Folr1* extraembryonic cell types is sufficient to produce severe embryonic phenotypes in a cell-non-autonomous manner (**Figure 6B, E, F, S6B**). Differential abundance analysis revealed similar patterns of cell-type depletion to our zygotic Δ*Folr1* mutants, including pronounced ablation of advanced trunk mesoderm and neural ectodermal cell types, with a relatively minor effect on endodermal cell types (**Figure S6D**). Notably, floor plate cells were the least depleted among neural cell types, consistent with our previous results and further supporting phenotypic concurrence between our zygotic and chimeric models.

**Figure 6.**
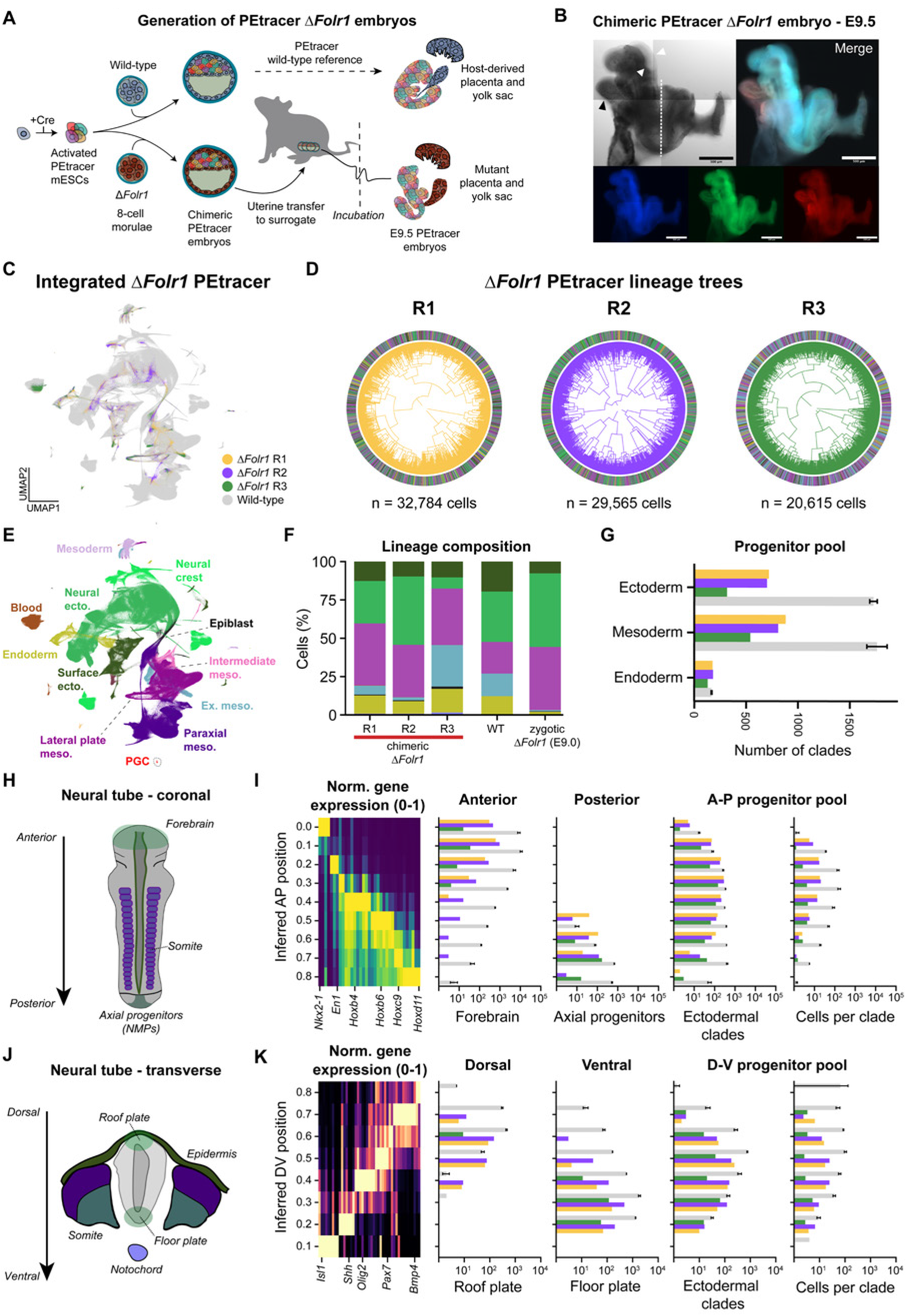
Chimeric PEtracer Δ*Folr1* embryos display distorted spatial organization. **A.** Schematic of chimeric mouse generation using Cre-activated PEtracer mESCs and zygotic *Folr1* knockout embryos. **B.** Representative whole-mount image of E9.5 chimeric Δ*Folr1* embryo morphology in sagittal view. White arrows denote the neural folds; a black arrow denotes the heart tube; a white dashed line highlights the underdeveloped caudal embryo. Fluorescent imaging highlights the pegRNA array (BFP, blue), PEmax (GFP, green), and lineage-tracing cassettes (mCherry, red). Scale bars: 500 μm. **C,E.** UMAP visualization of scRNA-seq data for three chimeric Δ*Folr1* embryo replicates, integrated with an E7.5-E10.0 PEtracer reference for wild-type mouse embryogenesis, colored by mutant replicate (**C**) and developmental lineage (**E**). **D.** Reconstructed lineage trees for 3 comprehensively sampled chimeric Δ*Folr1* whole-embryo replicates. Outer ring colors represent lineage assignments. R: replicate. **F.** Mean composition bar plots excluding blood for zygotic (E9.0) and chimeric (E9.5) Δ*Folr1* embryos, as well as the PEtracer reference (WT, E9.5). **G.** Number of fate-restricted clades per germ layer, colored by embryo replicate as in (**D**). Gray bars denote the reference mean at E9.5. **H,J.** Schematic of the developing neural tube in coronal (**H**) and transverse (**J**) view, highlighting the anterior-posterior (A-P) and dorsal-ventral (D-V) body axes, respectively. **I,K.** Distribution of cells and ectoderm-restricted clades by inferred spatial position along the anterior-posterior (**I**) and dorsal-ventral (**K**) body axes, with gray bars indicating reference and colored bars indicating Δ*Folr1* embryos, as in (**D**). <u>Left:</u> normalized expression (0–1) of genes used to infer A-P and D-V position, <u>center</u>: Log_10_ number of cells for curated cell types, <u>right:</u> Log_10_ number of fate-restricted clades and mean number of cells per clade. Error bars in **G,I,K** indicate the 95% confidence interval.

To examine the relative effects of folate depletion on embryo growth and differentiation, we reconstructed embryo-scale lineage trees for Δ*Folr1* chimeric embryos and systematically investigated progenitor dynamics throughout gastrulation and early organogenesis. Branch length estimates suggest that cell count in ΔFolr1 consistently diverges from wild type at approximately E4.0; however, because prime editing is sensitive to the concentration of free nucleotides^110^ which may be affected by folate depletion, our branch length estimates may be biased (**Figure S6C**). To mitigate this potential bias, we focused our analysis on the lineage architecture, including the number of fate-restricted clades and their relative output, rather than the timing of internal nodes. Quantifying the number of fate-restricted clades for each germ layer revealed that Δ*Folr1* PEtracer embryos exhibit substantial depletion of ectodermal and mesodermal progenitor pools relative to wild type. This pattern conforms with a consistent loss of proliferative potential in these lineages across replicates, despite evidence of compositional heterogeneity between samples (**Figure 6F, G**). We observed no such effect in the slower-cycling endoderm, indicating that proliferation defects and depletion are linked and favor relatively non-proliferative tissues, as our previous findings in the developing neural tube suggest.

To assess whether folate depletion affected cell-fate specification, we performed a comprehensive analysis of relative ancestral linkage across cell types and lineages. This approach revealed that lineage relationships were largely stable across replicates, indicating that *Folr1* disruption does not strongly affect lineage relationships (**Figure S6F**). The main difference was reduced linkage between the mesoderm, epiblast, and neural ectoderm, likely driven by the smaller NMP pool in Δ*Folr1* embryos. We also observed consistent relationships at the cell-type level, although we note a mild decrease in linkage between same-lineage cell types compared with wild type, which may reflect the developmental delay observed in the Δ*Folr1* condition (**Figure S6E**). Nonetheless, our tree-based analysis confirms that this perturbation largely does not affect differentiation trajectories; morphological and compositional disparities instead likely reflect differential expansion of germ layers and cell types.

The morphology of chimeric Δ*Folr1* embryos, including an underdeveloped posterior and an unclosed neural groove, further suggested distortions in cell distributions along the AP and DV body axes. Analysis of ectodermal progenitor pools along the AP axis revealed a marked depletion of ectoderm-restricted clades in the posterior neural tube and a parallel loss of proliferative output from these clades (measured as cells per clade) compared to wild-type, consistent with a disruption of posterior elongation (**Figure 6H, I**). Dorsal-associated clades were similarly affected, as Δ*Folr1* chimeric embryos displayed a pronounced preference for ventral identity, in agreement with the distorted relative abundance of the floor plate previously described in zygotic *Folr1* mutants (**Figure 6J, K**). Overall, these results strongly indicate that folate availability differentially affects growth in a germ layer-dependent manner, preferentially disrupting expanding tissues with high proliferation rates during primary neurulation, such as the elongating tailbud and dorsal neural tube. This effect further disrupts DV patterning because it spares slower-cycling “organizing” structures such as the floor plate, resulting in permanent morphological defects.

## Discussion

In this study, we explored the lineage-specific requirement of a ubiquitous, well-characterized metabolic pathway – folate-dependent OCM – within the dynamic context of gastrulation and early organogenesis. By characterizing metabolic states associated with early mouse development, we show that we can readily link transitions within well-defined biochemical pathways to distinct developmental events and spatial positions, with resolution otherwise unavailable through direct single-cell metabolic measurements. Through this approach, we identified a particularly sensitive metabolic transition specific to differentiation from pluripotent epiblast to neural ectoderm and associated with a high demand for *de novo* purine biosynthesis in a relatively hypoxic environment. This transition effectively unifies the morphological sensitivity of the neural tube with the unique metabolic requirements of neural differentiation, allowing us to explain seemingly lineage-specific defects in *Folr1*-deficient embryos by detailing the upstream metabolic drivers and downstream morphogenic consequences within the same analytical framework.

Our results suggest that the principal metabolic effect of severe folate deprivation lies in the 10-formylTHF-dependent steps connecting energy and purine metabolism, which preferentially affect actively expanding tissues such as the neural ectoderm during a brief window of animal development. Without abundant folate, we observe a breakdown in this connection, characterized by upregulation of downstream, folate-independent reactions in purine metabolism, presumably to support proliferation during a critical period of simultaneous growth and morphogenesis. While other functions of folate metabolism, such as DNA and histone methylation, affect cell fate specification in a lineage-specific manner^111,112^, we believe these effects are more subtle and downstream of the primary insult of nucleotide depletion, particularly given that our data show minimal differentiation effects along the neural trajectory. Further, our analyses indicate that purine metabolism defects in *Folr1* precede primary neurulation, suggesting an effect upstream of differentiation. The observation that lineage-specific proliferation defects then feed forward into permanent changes in morphology is also interesting, particularly the extent to which metabolic aspects of cell state correlate with the resulting over- or underabundance of neural cell types along the DV axis. Our PEtracer-based fate mapping confirms this observation, including lineage-specific effects on proliferative output that match the etiological understanding of folate deprivation in NTDs, including a window of vulnerability restricted to the period of axial elongation and neurulation and an increased incidence of spina bifida compared to anencephaly^113,114^.

Folate-dependent OCM is central to many facets of cellular physiology, including maintenance of redox homeostasis, homocysteine clearance, methyl donor availability, and, of course, support for cellular proliferation and transcription through nucleotide biosynthesis^16^. However, how these functions interact with tissue specification during early embryogenesis, and particularly how they produced lineage-specific defects in the formation of the neural tube, was less clear. Using MeRN, we extracted metabolic information from the transcriptomes of dozens of whole mouse embryos without bias. This insight revealed both a critical transitional period in neural lineage establishment and a breakdown in the metabolic activity that enables this transition following disruption of folate metabolism, thereby linking a ubiquitous metabolite to a neural-associated phenotype.

The ability to describe biological systems at the single-cell level has been a defining aspect of the 21st-century, driven by increasingly large and complex datasets from countless species across their life courses and from large cohorts of human patients. The rise of these technologies has catalyzed the development of new tools that integrate and interpret these data, including analytical approaches that can estimate physical or cellular parameters from what is effectively genomic data. As a technique, MeRN demonstrates the power of deep learning, particularly by using metabolic topology as an applied inductive bias, to interrogate outstanding questions in biology with widely available data structures. This framework is likely to have broad potential, particularly in areas where the relationship between metabolism and cell fate directly impacts tissue organization or disease-associated dysregulation^40,115^. Alongside other transcription-driven metabolic analysis methods^36–40^, MeRN represents a substantial step forward in probing these relationships, enabling new avenues for unbiased, scalable investigation using readily accessible platforms and tools.

Our dissection of folate metabolism in the developing embryo was an excellent candidate for this approach, particularly given its historical epidemiological link to neural tube defects and nutritional fortification. The ability to investigate rapid metabolic changes occurring within the twelve hours that bridge gastrulation and early neural specification, which require exceptional coordination between growth and differentiation, was indispensable to our analyses and interpretation. Translationally, these results also support nucleotide precursor-based supplementation therapy as a more specific route to prevent NTDs during gestation^116^ and further support glycerolipids and arachidonic acid as early biomarkers for developing congenital abnormalities^109^.

### Limitations of the study

While our application of this framework to multiple independently generated datasets of mouse embryogenesis suggests broadly consistent patterns, it remains to be seen whether the metabolic states identified here are generalizable to other systems and models, and more importantly, whether they provide similarly effective descriptions of metabolic dynamics, diversity, differential activity, and susceptibilities in response to experimental conditions. MeRN relies on a curated set of metabolic genes and currently does not account for transporters and receptors, such as FOLR1 or the more ubiquitous reduced folate carrier (RFC), which may modulate responses to metabolic insults. The model also relies on transcript-level data, and as such, may not accurately reflect the full repertoire of metabolic flux. Second, while our genetic model mitigates confounding effects of altered maternal diet, it indirectly mediates folate availability; further studies should compare whole-embryo and extraembryonic-localized disruption to identify distinguishing traits between these systems. Additionally, we do not directly target purine biosynthesis through genetic ablation of enzymes catalyzing the folate-dependent reactions highlighted by *Gart* and *Atic*, which have underdescribed phenotypes characterized primarily by preweaning lethality and decreased proliferation^117,118^. Third, our expanded array of mutants was temporally restricted to E8.5, prohibiting analysis of altered metabolism dynamics throughout development. Fourth, the PEtracer system is sensitive to nucleotide availability; therefore, our branch-length estimates should be considered as internally controlled, not as absolute values. Finally, we focused our analyses on neural effects of folate uptake restriction, but other tissue morphogenesis may be underexplored, such as in cardiogenesis and extraembryonic tissues where *Folr1* is primarily expressed.

## Acknowledgements

We thank the members of the Smith Lab, past and present, for their support in the development of this manuscript. We thank S. Reilly, M. Chan, and B. Sozen for their continued guidance and discussions during manuscript assembly. We also thank H. Kretzmer, R. Tornisiello, M. Morales, and T. LaMoia for helpful discussions and support.

## Author contributions

ND, DPL, AW, and ZDS conceptualized and designed the embryogenesis analyses using MeRN. MW, JV, and ZDS generated and collected mutant embryo cohorts and scRNA-seq libraries; ND, YK, and MW prepared the scRNA-seq data used for these analyses. DPL and AC implemented MeRN; ND and DPL analyzed and interpreted its outputs. WNC, JV, TH, GG, and LWK generated and analyzed PEtracer embryos and their scRNA-seq data; WNC, LWK, DPL, and ND interpreted results. MW, JV, and KS performed imaging-based validation of embryo samples. ND, DPL, and WNC interpreted analytical outputs under the supervision of and with support from TA, JSW, LWK, AW, and ZDS. ND, DPL, AW, and ZDS wrote the manuscript, with input from all authors. All authors reviewed and approved the final manuscript.

## Funding

ZDS is supported by the National Institutes of Health (NIH) Director’s New Innovator Award (DP2HD108774), the Mathers Foundation, the Chen Innovation Award, and the Max Planck Society. Y.K. is supported by an EMBO postdoctoral fellowship (ALTF 315-2024). DL, AC, and AW were supported by a research grant from the Shurl and Kay Curci Foundation. LWK is supported by the Helen Hay Whitney/HHMI fellowship and Eunice Kennedy Shriver NICHD Pathway to Independence Award NIH K99HD118574. JSW is supported by the Howard Hughes Medical Institute, the National Institutes of Health (NIH) Centers of Excellence in Genomic Science (RM1HG009490), the Chan Zuckerberg Initiative 2024-346405 (5022), and the Whitehead Innovation Initiative.

## Declaration of interests

ZDS is an academic cofounder and scientific advisor to Harbinger Health. WNC consults for Merck Pharmaceuticals. JSW declares outside interests in 5 AM Venture, Amgen, nChroma Bio, DEM Biosciences, KSQ Therapeutics, Maze Therapeutics, Tenaya Therapeutics, Tessera Therapeutics, Thermo Fisher Scientific, and Xaira Therapeutics.

## Declaration of generative AI and AI-assisted technologies in the writing process

The authors wrote the manuscript, and AI was used to improve grammar and clarity. For figure legends and methods, agentic AI with access to the codebase was used to verify that the textual description fully aligns with the codebase. All authors reviewed the final manuscript and take full responsibility for its contents.

## RESOURCE AVAILABILITY

### Lead Contact

Please direct information and resource requests to the lead contact, Zachary D. Smith, who will fulfill them.

### Materials Availability

Reagents generated in this study are available upon request from the lead contact, and other materials are commercially available.

### Data and Code Availability

All code and data used in this work will be made available at the time of publication.

## METHOD DETAILS

### Mouse strains and husbandry

Experimental mice were housed and maintained in accordance with ethical guidelines approved by the Yale University Institutional Animal Care and Use Committee (IACUC Protocol #2026-20357) and the LaGeSo Berlin Institutional Animal Care and Use Committee (Protocol #G0098/23). All procedures adhered to governmental and public health service regulations. Mice were maintained in specific pathogen-free facilities under a 12-hour light/dark cycle (lights on 7:00-19:00). This study utilized B6/CAST F1 male mice generated in-house by breeding C57BL/6J strain female mice with CAST/EiJ strain males as sperm donors for ICSI alongside 6-12-week-old B6D2F1/J (BDF1, Jackson Laboratory or Janvier) female mice to provide oocytes. 25-35 g female CD-1 mice were purchased from Charles River Laboratory or in-house colonies for use as pseudopregnant surrogates, and Swiss-Webster or CD-1 strain vasectomized males (9 weeks old) were purchased from Taconic Biosciences or in-house colonies.

### Embryo generation, collection, and staging

Mouse embryos were generated using previously described methods combining intracytoplasmic sperm injection (ICSI) and CRISPR-based genome editing^34,41,42,79,80^. Briefly, BDF1 strain females (Charles River Laboratory, Janvier) were superovulated by serial injection of 5IU of CARD HyperOva FD (Cat. #KYD-015-EX, Cosmos Bio) and 5 IU of human chorionic gonadotropin (Cat. #HOR-250, Prospect Protein Specialists) separated by 44-48 hours. MII stage oocytes were isolated the following morning, and cumulus cells were removed using M2 media with hyaluronidase (Cat. #MR-051-F, Millipore Sigma), followed by stable culture in pre-gassed KSOM Advanced media (Cat. #MR-101-D, Millipore Sigma) overlaid with pre-tested mineral oil. Thawed CAST sperm were injected into oocytes using piezo-assisted intracytoplasmic sperm injection (ICSI) with 6 µM I.D. glass injection needles using an Eppendorf Transferman micromanipulation system and a Hamilton Thorne I.R. laser. After 4–6 hours following sperm injection, Pronuclear stage 2–3 zygotes were injected with a cocktail of Cas9 and sgRNA mRNAs (200 ng/µL Cas9, 100 ng/µL sgRNA pool) and cultured to the blastocyst stage before uterine transfer into pseudopregnant CD-1 strain female donors.

### sgRNA design and synthesis

sgRNAs were designed by examining the gene of interest in the UCSC Genome Browser (mm10 assembly) and selecting target exons shared across all reported isoforms. Individual exons were selected to maximize coverage across the gene of interest with the fewest sgRNAs possible, typically 3 to 5, and individual guides were designed using the Integrated DNA Technologies CRISPR Cas9 guide RNA design checker, carefully inspecting potential off-target effects and selecting protospacer candidates with good base complexity and few, if any, potential off-targets with fewer than three mismatches to the closest off-target locus. Prospective sgRNAs were then counter-screened for the presence of common SNPs in the UCSC reference to prevent strain-specific targeting. Once selected, individual sgRNAs were generated via PCR using T7-promoter-containing primers and 2x Phusion Master Mix (Cat. # F531L, Thermo Scientific) to create templates for in vitro transcription. Individual T7 guide templates were pooled in an equimolar ratio, followed by *in vitro* transcription using the HiScribe T7 High Yield RNA Synthesis Kit (Cat. #E2050S, NEB). We purified IVT reactions using the Monarch RNA Clean-up Kit (Cat. #T2040L, NEB) and resuspended them in Injection Buffer (5 mM Tris buffer, 0.1 mM EDTA, pH 7.4).

### Imaging-based validation of embryo phenotypes

For whole-mount DAPI and phalloidin staining, embryos were fixed in 4% PFA overnight at 4°C, then washed and stored in 1X PBS. Embryos were placed in PBS + 0.2% Triton X-100 (PBS-T) containing DAPI (Sigma, D9542-5MG) and TRITC-Phalloidin (Millipore Sigma, P1951-0.1MG) and incubated with gentle rocking for 4–6 hours at room temperature. For whole-mount imaging, samples were washed 3 x 10 minutes with PBS-T and incubated in Ce3D media (22% N-methylacetamide [Sigma, M26305-100G], 86% Histodenz [Sigma, D2158-100G]) overnight at room temperature for optical clearing. We transferred cleared embryos to a 35 mm glass-bottom dish for confocal imaging (MatTek, P35G-1.5-20-C). Images were acquired on an inverted Leica Stellaris 5 confocal laser scanning microscope using Diode 405, a white light laser, and *LAS-X* software (v4.6.1; Leica) using an HC FLUOTAR L VISIR 25x/0.95 water objective.

For anterior-posterior axis-based analysis of the neural tube, fixed embryos were stained with Foxa2 (1:500; Rabbit, #8186S, CST), Sox2 (1:800, rat, #14-9811-82, Invitrogen) primary and Donkey anti-Rat IgG (1:1000, Alexa Flur 488, #A48269, Invitrogen), Donkey anti-Rabbit IgG (1:1000, Alexa Flur 555, #A31572, Invitrogen) secondary antibodies prior to light sheet imaging on the Zeiss Z1 Lightsheet microscope. After imaging, we cryosectioned embryos for high-resolution analysis at set points along the body axis and imaged them with a Zeiss LSM880.

### scRNA-seq generation and pre-processing

#### Single-Cell RNA-seq Library Preparation and Sequencing

We isolated whole mutant embryos from surrogate female mice at the desired time point of interest and carefully cleaned them of maternal tissue. Briefly, individual deciduae were collected in 1x PBS, then serially washed through several drops before collecting ten or more embryos per time point for single-cell analysis. Typically, embryos were collected from two or more surrogates to reduce batch-level variation in developmental progression, and embryos did not undergo further screening to avoid observational bias. Transferred embryos were then transferred into a 200 µL drop of 0.2% BSA/PBS and dissociated into single cells by replacing the 0.2% BSA/PBS with TrypLE (Cat. #12604013, Invitrogen), followed by mechanical dissociation using a P200 pipette every 5 minutes until cells reached a clear single-cell suspension. Cells were then serially quenched and resuspended in 0.2% BSA/PBS in a LoBind Eppendorf tube (Cat. #22431021, Eppendorf) and filtered using a Scienceware Flowmi cell strainer, 40 µm (Cat. #10032-802, VWR). After at least three washes, cells were resuspended in a final volume of 100 µL and manually counted on a hemocytometer. Cells were then normalized to target a recovery of 12,000 cells prior to microfluidics-based single-cell profiling using the Chromium Single Cell NextGem 3’ v3 system (10x Genomics). For early time points, we ran a single reaction; for embryonic stages after E8.0, we ran two reactions. Single-cell libraries were prepared following the protocol with a cDNA and PCR indexing amplification cycle of 10 and examined on a D5000 and D1000 TapeStation System, respectively (Agilent Technologies). Libraries were sequenced to a minimum of 350 million paired-end reads according to the manufacturer’s protocol using either the Illumina NovaSeq or Element Biosciences AVITI systems.

### scRNA-seq data preprocessing

We applied a previously described preprocessing strategy^34,41^ to all mutant embryo data, including sequence alignment, quality control, ambient RNA removal, and doublet identification. Briefly, we first processed sequencing FASTQ files with 10x Genomics *CellRanger*^119^ (v6.0.2) for genome alignment and gene expression quantification. We then passed the unfiltered gene expression matrices to *CellBender*^120^ (v0.3.0) to remove ambient RNA contamination from lysed cells in the 10x Chromium input suspension. We converted the output h5ad file to a Seurat object using *SeuratDisk* (v0.0.0.9021) and *Seurat*^121^ (v5.1.0). To remove low-quality cells, we calculated the median absolute deviation (MAD) of log-transformed UMI counts of single cells and set the minimum UMI count threshold as:

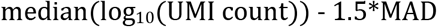

If the back-transformed UMI threshold was lower than 2000, we manually set it to 2000. Next, we removed doublet cell barcodes using *Scrublet*^122^ (v0.2.3), which assigns a doublet score to each cell barcode by simulating synthetic doublets and computing the transcriptional similarity of each observed cell barcode to the simulated doublet population. We set the expected doublet rate to 7.5%. For doublet simulation, we identified highly variable genes using a minimum gene variability percentile of 85% and used the first 20 principal components for transcriptome embedding. We visually inspected the doublet score distribution and set score thresholds at local minima between peaks in bimodal distributions, with a default threshold of 0.25 for unimodal or mixed distributions.

### Embryo demultiplexing

We performed embryo demultiplexing using the workflow described by Grosswendt et al.^34^. Briefly, the cDNA reads in Read 2 were aligned to the mouse reference genome *mm10* using *STAR*^123^ (v2.5.3), and the alignment BAM file was split into B6 (reference) and CAST (alternative) using *SNPsplit*^124^ (v0.3.4). We then counted the read counts assigned to two genotypes for each chromosome in a single cell and transformed them into a cell-by-chromosome allelic read-fraction matrix. Using only autosomes, we first filtered cells with at least 1,000 SNP-informative read counts (i.e., reads overlapping known B6/CAST SNP positions). Then, we performed k-means clustering on the allele fraction matrix with k equal to the number of pooled embryos. For each k-means cluster, we calculated the Mahalanobis distance from each cell to the cluster centroid and filtered out cells with a distance greater than one MAD above the median, retaining only confidently demultiplexed cells.

To visualize the demultiplexing result, we applied *UMAP*^125^ to the cell-by-chromosome allelic read-fraction matrix, using 15 neighbors to project into 2 dimensions with the cosine distance metric and a minimum distance of 0.1 with the R *uwot* package (v0.1.11). When we split a single-cell suspension across two GEM wells, we merged data from both wells before demultiplexing. When we loaded each suspension into an independent GEM well, we applied the pipeline separately to each well.

### Embryo rank ordering

To order embryos along an inferred developmental trajectory, we applied a custom embryo-ranking pipeline^42,73^. Extra-embryonic ectoderm and extra-embryonic endoderm were excluded from this analysis. We first ranked wild-type and Δ*Folr1* embryos separately by transcriptional similarity using per-embryo pseudobulk gene expression profiles (summed raw counts). We normalized these pseudobulk data and identified the top 3,000 most variable genes for scaling and PCA (30 principal components). We computed a Euclidean distance matrix from the PCA embeddings and converted it to a pairwise similarity matrix by taking the inverse of each distance value, adding a pseudocount of 0.0001 to avoid division by zero.

We then ordered embryos to maximize local transcriptional similarity using a custom iterative bubble-sort algorithm, as in Wang et al.^42^. Briefly, we compared adjacent embryo pairs by computing the difference in their summed similarity scores to all non-adjacent embryos; if swapping a pair increased the ordering’s total local similarity, we performed the swap. We repeated this procedure iteratively until no further swaps improved the ordering or until we reached a maximum of 500 iterations. We used the resulting order as the developmental ranking of each embryo within each condition, as shown in **Figures 3E and S2B, D**.

### Cell-level pseudotime assignment

To assign continuous developmental pseudotime to individual cells, we leveraged our previous embryo rankings and *Seurat*’s anchor-based label-transfer framework to assign cells from either the wild-type or Δ*Folr1* datasets to a wild-type embryo. Specifically, we identified transfer anchors between the wild-type reference and each query dataset (wild-type self-reference and Δ*Folr1*) using the first 30 principal components of the wild-type (reference) PCA embedding, calculated from log-normalized, scaled expression of the top 3000 most highly variable genes. For each cell, label transfer produced a vector of prediction scores across all 72 wild-type embryos, reflecting each cell’s relative transcriptional similarity to each wild-type embryo, normalized to sum to 1.

Developmental pseudotime was then computed as a weighted linear combination of embryo ranks, where each cell’s prediction score vector was multiplied by a rank index vector (1 through *n*, where *n* = 72 is the total number of wild-type embryos) corresponding to the rank order described above. The resulting scalar value for each cell reflects its weighted position along the inferred developmental trajectory, with higher values indicating a more advanced developmental stage. We assigned this pseudotime score to all cells in both the wild-type and Δ*Folr1* datasets.

### Cell type label transfer

To assess the transcriptional identity of Δ*Folr1* cells relative to the pre-annotated wild-type reference, we performed label transfer using Seurat’s anchor-based framework. We identified transfer anchors as described above. We then predicted cell type labels for each Δ*Folr1* cell, producing a per-class prediction score (summing to 1 across classes) and a maximum prediction score reflecting the confidence of the best-matching cell type assignment.

To establish a baseline for label prediction, we additionally performed label transfer using the wild-type dataset as both reference and query. The resulting self-reference maximum prediction score distribution served as an upper-bound reference distribution against which we compared Δ*Folr1* scores using two complementary metrics. We applied a two-sided Kolmogorov-Smirnov (KS) test to quantify the maximum vertical difference between the empirical cumulative distribution functions (ECDFs) of wild-type and Δ*Folr1* prediction scores, measuring the greatest local divergence between distributions. We also computed the Wasserstein distance to quantify the total area between ECDFs, capturing the overall magnitude of the distributional shift. KS tests were performed using the base R *stats* package^126^ (v4.2.3), and Wasserstein distances were computed using the *transport* package (v0.15-4).

To examine lineage-specific prediction confidence for each cell type, we computed the mean maximum prediction score per cell type per condition, excluding germ cells due to their low abundance. We used these scores to identify cell populations with notably reduced mean prediction scores relative to the self-reference baseline.

#### Applying the Metabolic Representation Network (MeRN)

We applied the MeRN model, latent space inference, reaction activity inference, and DDP construction as described in the accompanying manuscript by Lewinsohn et al. In total, we applied MeRN to three data subsets: the full wild-type scRNA-seq data and two data subsets excluding extra-embryonic endoderm, extra-embryonic ectoderm, and blood cell lineages for both wild-type and Δ*Folr1* embryo data. Here, we refer to these as the *full* model and the wild-type or Δ*Folr1 embryonic* models, respectively. Below are training details, the number of model replicates, and thresholds. We used MeRN v1.0.0.

### MeRN model training

For all three data subsets, we trained 10 MeRN model replicates. Each replicate was trained with latent dimensions *z* = 25 (metabolic), *u* = 15 (non-metabolic), Kullback-Leibler (KL) divergence weights *β_cell_* = 0.001, and *β_ggrap_*_ℎ_ = 0.1, with loss-balancing hyperparameters *λ*_data_ = 1 and *λ*_graph_ = 0.2. We performed model optimization using the root mean square propagation (RMSprop) optimizer with a learning rate of 2 × 10^−3^ and a batch size of 128. We split cells into training and validation sets using a 90/10 split. We used the total validation loss for early stopping and selected the model with the lowest validation loss for downstream analyses.

### Inference of single-cell reaction and KEGG pathway activity

To infer metabolic reaction activity in single-cells, for each of the 10 MeRN model replicates from each data subset, we randomly sampled 8 reaction activity estimates and took the average, resulting in a cell-by-reaction activity matrix for each model replicate. We then averaged the reaction activity matrices across all 10 replicates for each data subset. Further details and rationale are provided in Lewinsohn et al.

To calculate KEGG pathway activity scores, we averaged the reaction scores for component reactions (**Table 2**). We filtered pathways to those with validated KEGG *Mus musculus* orthologs (**Table 3**) and removed pathways with fewer than 5 component reactions present in the MeRN inference, as well as those denoting overarching global and overview maps (i.e., those whose ID began with **rn01**).

### Prediction of embeddings and reaction activity in mutant embryos

To predict latent space embeddings for mutant cells, we passed their gene expression profiles through the encoder of the first MeRN model replicate trained on the wild-type embryonic data subset. To calculate reaction activities for the mutant data, we performed 8 forward passes through each of the 10 MeRN replicates trained on wild-type embryonic data, then averaged reaction activity across samples and replicates for downstream analyses. We generated two-dimensional representations using the *SCANPY*^127^ (v1.11.5) implementation of the UMAP^125^ algorithm with default parameters, using either metabolic or non-metabolic latent-space embeddings for each visualization.

### Metabolic state annotation

To identify metabolic states, we applied Leiden^128^ clustering to the metabolic latent space derived from the first MeRN model replicate trained on the embryonic wild-type data subset, using an empirically determined resolution of 0.4. We defined metabolic state annotations based on community size, with MS0 as the largest community and MS5 as the smallest.

### Differential pathway activity between MS2 and MS5

To identify pathways differentially active between MS2 and MS5 in wild-type samples, we compared embryo-level pseudobulk KEGG pathway activity scores across these two states. Only embryos contributing at least 50 cells to either MS2 or MS5 were retained for this analysis. For each qualifying embryo, we averaged pathway scores (derived above) across all cells within each metabolic state, producing one pseudobulk score per embryo, per state, per pathway.

For each pathway, we quantified effect sizes using Cohen’s d with a pooled standard deviation estimator, with positive values indicating higher activity in MS5 relative to MS2.

### Metabolic state prediction in mutant embryos

To assign metabolic states to mutant cells, for each dataset, we first calculated their 500 nearest neighbors in the shared metabolic embedding derived above, where the neighbor pool includes both wild-type and mutant cells. We then assigned the metabolic state label held by the majority of each cell’s 10 closest wild-type neighbors. If no state held a majority, or if fewer than 10 wild-type cells were among the 500 nearest neighbors, we did not assign a metabolic state label to the mutant cell.

### Spatial data analysis and metabolic state prediction

We conducted all spatial transcriptomics analyses on the seqFISH data produced by Harland et al.75 The preprocessed data are publicly available (**see Data and Code Availability**).

To predict metabolic states in the spatial data, we first identified marker genes for each state using a cell-level one-versus-all Wilcoxon test on log-normalized expression profiles from the wild-type scRNA-seq data. We then constructed a metabolic state gene set from the top 100 upregulated genes for each state, ranked by standardized Wilcoxon rank-sum test statistics.

We intersected each gene set with the 351-gene seqFISH probe target panel used by Harland et al., resulting in 13 available genes for MS0, 19 for MS1, 16 for MS2, 7 for MS3, 14 for MS4, and 8 for MS5. We then used SCANPY’s *score_gene* function to calculate scores for each gene set, standardized these scores across all cells from all combined replicates (*n =* 7 embryos), and assigned metabolic states based on the highest standardized gene set score.

### Data-driven pathway (DDP) construction and application

To cluster reactions into functional metabolic sub-units, we constructed DDPs as described in Lewinsohn et al. Briefly, we first calculated pairwise Spearman correlations between reactions across wild-type MS2 cells. We selected this metabolic state because it is abundant in both wild-type and ΔFolr1 samples and represents the common state before gastrulation (E6.5-E7.5). To group reactions into DDPs, we applied graph-constrained agglomerative clustering using metabolic topology connectivity (i.e., adjacency between reactions as defined by the graph representation of metabolism used by MeRN) and Spearman correlation between reactions. We applied a minimum correlation threshold of *ρ* = 0.7 and a minimum DDP size of 3 reactions.

To calculate DDP scores per cell, we averaged the reaction scores of component reactions per DDP, as described for KEGG pathways (**Table 2**).

### Differential DDP activity between wild-type and Δ*Folr1*

To identify DDPs differentially active between wild-type and Δ*Folr1* cells in MS2, we applied the same pseudobulk differential activity framework described above, using DDP scores in place of KEGG pathway scores and restricting the analysis to MS2 cells. We excluded embryos contributing fewer than 50 MS2 cells. Cohen’s d positive values indicate higher activity in Δ*Folr1* relative to wild-type.

### Analysis of relative DDP activity across mutant embryos

To characterize global relationships between embryos across genotypes, we applied the same per-embryo pseudobulk DDP aggregation described above and submitted the resulting embryo-by-DDP matrix to PCA using centered and scaled scores.

To quantify how much each mutant embryo’s DDP activity profile deviated from wild-type, we computed two complementary measures using wild-type embryos as a reference. First, we z-scored each mutant embryo’s mean DDP activity relative to the wild-type distribution (mean and standard deviation computed across all wild-type embryos). Second, we computed the Euclidean distance of each mutant embryo’s mean DDP activity vector from the wild-type centroid, providing a single scalar measure of overall metabolic divergence from the unperturbed baseline.

### Reaction co-activity analysis for transformylation reactions

To assess the co-activity of two folate-dependent purine metabolism reactions across development and between genotypes, we computed per-embryo Spearman correlations between the inferred activity scores of GAR transformylase (**R04325/R06974**) and AICAR transformylase (**R04560/R06975**). Reaction activity scores for each condition were independently derived from the MeRN models trained on the wild-type embryonic data subset for wild-type cells, and the Δ*Folr1* embryonic data subset for Δ*Folr1* cells, as described above, and combined into a single analysis dataset. We excluded cells that could not be assigned to a majority metabolic state, as well as cells from E6.5 embryos, for which no time-matched Δ*Folr1* data were available. Only embryos contributing at least 20 cells were retained.

For each qualifying embryo, we computed the Spearman correlation between the two reaction activity scores across all retained cells. We then tested for differences in per-embryo correlation coefficients between wild-type and Δ*Folr1* embryos at each developmental time point using the hypothesis-testing framework described in **General analysis conventions**.

### Metabolic Pathway Visualizations

To visualize metabolic reaction connectivity in context, we used the *custom_pathway_plot* function from *MeRN* (v1.0.0) in *Python* (v3.10.20). We constructed these graph layouts based on the KEGG^44^ reaction graph used by MeRN, representing metabolites as nodes and reactions as edges, and including only edges between metabolites that were substrate-product pairs in an inferred reaction. Metabolite node coordinates were selected using the *graphviz_layout* function from *NetworkX* (v3.4.2). We note that metabolic pathway visualizations approximate, and thus imperfectly represent, the underlying metabolic topology.

### scRNA-seq analysis

#### General analysis conventions

All significance tests were performed with a two-sided Mann-Whitney U test, unless otherwise noted, and p-values were adjusted with the Benjamini-Hochberg procedure for multiple hypotheses when appropriate. Results with adjusted p-values smaller than 0.05 were considered significant, unless otherwise noted.

Of the 94,596 cells in the wild-type dataset, 20,381 (21.5%) failed sample-level demultiplexing and were excluded from all pseudobulk analyses.

### Inference of hypoxia scores per embryo

To infer the relative timing of hypoxic conditions over the developmental trajectory, we computed a composite hypoxia score per embryo rank (derived above) using the mouse MSigDB Hallmark Hypoxia^74^ gene set. We filtered genes in the set to those present in the wild-type scRNA-seq dataset and expressed above a minimum threshold (mean log-normalized count across all cells > 0.01 and detected in at least 5% of cells), retaining 139 (70.0%) of the 198 genes in the set (**Table 1**).

For each retained gene, we averaged log-normalized values across wild-type cells within each embryo rank, irrespective of cell type, to generate a whole-embryo pseudobulk expression profile per rank. We then computed a composite hypoxia score per rank by averaging across all retained genes, and applied z-score normalization across all rank positions.

To visualize the hypoxia score across development, we plotted the z-scored values per embryo, showing individual embryos as points ordered by rank and overlaid on a loess-smoothed trend line, using default *ggplot2*129 (v4.0.1) parameters.

### Estimation of embryo cell counts

To estimate cell counts per embryo, we first calculated the proportion of cells in each scRNA-seq dataset attributable to each embryo. We then multiplied this per-embryo proportion by the product of the total cell count (obtained by manual hemocytometer counting during scRNA-seq library preparation) and the dilution factor (Table 4). To estimate cell counts per lineage, we applied a similar approach: we calculated the proportion of cells assigned to each lineage label (endoderm, mesoderm, non-neural ectoderm, and neural ectoderm) within each embryo, then multiplied by the estimated total cell count per embryo derived above.

To evaluate significance between wild-type and Δ*Folr1* embryo and lineage cell counts, we used a one-sided Mann-Whitney U test, testing the hypothesis that mutant cell counts were lower. Resulting p-values were adjusted with the Bonferroni correction.

### Cell cycle analysis with tricycle

We estimated cell cycle positions using the R *tricycle* package^94^ (v1.6.0), applied separately to the wild-type and Δ*Folr1* scRNA-seq data using the *Runtricycle* function from *SeuratWrappers* (v0.3.5), specifying the *species* parameter as “mouse” with otherwise defaults applied. Cell cycle phases were assigned based on position, where θ ∈ (0,2π), using the following thresholds: *S* (0.5π ≤ θ < π), *G2* (π ≤ θ < 1.5π), *M* (1.5π ≤ θ < 1.75π), and *G0/G1* (θ ≥ 1.75π or θ < 0.5π).

We assessed statistical differences between conditions in the neural ectoderm with Watson’s two-sample test of homogeneity^130,131^, a non-parametric test for the null hypothesis that two samples are drawn from the same circular probability distribution. This test is therefore appropriate for testing whether two sets of embryos are homogeneously distributed along the (circular) cell cycle axis. We used the *circular* package (v0.5-1) for this analysis and a permutation strategy to assess significance. Briefly, for each of the four cell classes (floor plate, neural crest and NMP, neurons, and neural epithelium, fore-, mid-, and hind-brain), we computed the Watson *U^2^* statistic between wild-type and Δ*Folr1* cells, then generated a null distribution of *U^2^* by randomly permuting condition labels in each sample (*B* = 999 permutations), recomputing the test statistic at each iteration, and computing an empirical p-value as:

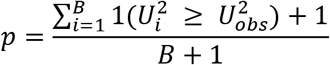

That is, the p-value is equal to the number of statistics drawn from the null distribution with values greater than or equal to the observed statistic, divided by the number of permutations. We also applied a Laplace correction (+1) to the numerator and denominator, and consequently bounded the lowest p-value at 0.001 for this number of permutations.

### Differential abundance analysis with Milo

We performed differential abundance analysis using Milo^87^ as implemented in the *R* package *miloR* (v1.6.0) to identify cell populations differentially represented between wild-type and Δ*Folr1* samples. The wild-type E6.5 time point was excluded from this analysis due to the lack of a time-matched sample in the Δ*Folr1* condition. We first merged the processed Seurat objects and subset these data to exclude extra-embryonic endoderm, extra-embryonic ectoderm, and blood cell types. We converted this object to a *SingleCellExperiment*^132^ object (v1.20.1) for cross-platform compatibility, then to a Milo object.

We performed two parallel Milo analyses, using k-nearest neighbor (kNN) graphs calculated with either metabolic (*d* = 25) or non-metabolic (*d* = 15) latent space embeddings. We defined transcriptional neighborhoods using the graph-refinement scheme (*prop* = 0.01). We iteratively adjusted *k* based on visual inspection of neighborhood-size distributions, targeting a median neighborhood size of 50–100 cells and arriving at *k* = 30 (metabolic) and *k* = 50 (non-metabolic). We counted cells per neighborhood, with each sequencing run as the sample-level replicate unit. Neighborhood distances were calculated using the same dimensions as each kNN graph. We tested differential abundance between wild-type and ΔFolr1 using a generalized linear model with experimental condition as the sole fixed effect and considered neighborhoods with a spatial FDR < 0.05 as statistically significant^133^. Neighborhoods calculated using the non-metabolic latent space graph were annotated by majority cell type and lineage labels, and those calculated with the metabolic latent space graph were annotated by majority metabolic state.

### Cell type network flow inference

To characterize cell type transitions during mouse development, we reconstructed a directed network of cell type flows for the wild-type reference previously computed by Kijima et al.41 and based on the framework first described by Mittnenzweig et al.73 from aggregated metacell-level flow data.

Briefly, we summed pre-computed forward flow matrices generated from leave-one-out cross-validation per embryo (*n* = 72 total flows) across all replicates to produce a single aggregated network flow graph, where each entry represents the total flow between a pair of metacells across the dataset. Metacell cell types were pre-annotated by Kijima et al.^41^

To obtain a directed, unidirectional representation of the net flow between metacell pairs, we computed the difference between the forward flow matrix and its transpose, retaining only edges with a net weight greater than 5. This step removes reciprocal flow and retains only the dominant, forward direction of transcriptional transition between each metacell pair. We then aggregated metacell-level net flows to the cell-type level. For each directed cell type pair, we summed net edge weights across all contributing metacell pairs and normalized by the number of metacells in the origin cell type, producing a mean net flow weight per transition. To ensure consistent directionality after cell-type-level aggregation, we repeated the forward-minus-reverse subtraction at the cell-type level. We retained only edges with positive net weight, removing bidirectional redundancy introduced by aggregation.

### Flow network graph construction and layout

The resulting directed edge list was used to construct a graph using the *igraph* (v2.1.3) and *tidygraph* (v1.3.1) frameworks. We annotated nodes with lineage identity for downstream filtering. To determine cell-type placement, we counted cells per embryo rank (derived in **Embryo rank ordering**) for each cell type and arranged them into a cell-type-by-rank matrix. We then normalized each column by the total number of cells across all cell types at that rank. Hence, each entry reflects the relative proportion of a given cell type at a given rank, controlling for differences in total cell numbers across embryos. We defined the representative rank for each cell type as the rank with the highest normalized column proportion, indicating the embryo rank at which that cell type was most relatively enriched compared with other cell types, and used this modal rank to position cell type nodes along the x-axis of the network layout.

We subset the network to exclude extra-embryonic endoderm, extra-embryonic ectoderm, and blood lineages. We manually arranged nodes in a two-dimensional layout, ordering cell types vertically according to a fixed lineage order (from extra-embryonic mesoderm to endoderm, with the epiblast centered), and visualized them using *ggraph* (v2.2.1). We determined timepoint boundaries based on the maximum rank within each developmental stage (E6.5-E8.5), indicated as vertical reference lines.

### Calculation of cell-type-level pathway activity in the flow network

To overlay metabolic pathway activity onto the flow network, we used the per-cell KEGG pathway activity scores for glutathione metabolism (**rn00480**) and glycolysis and gluconeogenesis (**rn00010**) derived above. We removed cells from lineages excluded from the flow network and averaged pathway scores per embryo-cell type combination (pseudobulk), retaining only those with at least 20 cells. We z-score-normalized pathway activity scores across all embryo-cell type observations within each pathway and then averaged these scores across embryos per cell type. We then assigned the resulting per-cell-type mean scaled pathway activity scores to network nodes and visualized them in the graph layout.

### PEtracer chimeric embryo generation and analysis

#### Generation of PEtracer mESC-derived chimeric embryos

We conducted PEtracer mESC-based recording as previously described by Colgan et al.^43^ but using Folr1 mutant precompaction 8-cell-stage embryos instead of diploid CD-1 strain donor embryos used to generate the reference data. Briefly, 1–2 million PEtracer cells were nucleofected with 10 µg pCAG-Cre plasmid (Addgene plasmid #13775) using the P3 Primary Cell 4D-Nucleofector® X Kit L (Lonza, V4XP-3024) and the CG-104 program on a Lonza 4D-Nucleofector X system, following the manufacturer’s instructions and as previously described^134,135^. At 24 hours post-nucleofection, cells were trypsinized and sorted for mCherry and GFP double-positive populations using a BD FACSAria™ II Cell Sorter (BD Biosciences) operated with BD *FACSDiva* software (v8.0.1), including identical gating parameters as used previously. GFP-positive cells were defined relative to non-transfected control cells to establish background fluorescence, and the top approximately 80% of GFP-expressing cells within the mCherry+/GFP+ double-positive population were sorted and maintained on ice until injection. Subsequently, 6–8 healthy PEtracer mESCs were injected into pre-compaction 4- to 8-cell stage embryos, cultured for an additional 24 hours, and transferred into the uterine horns of pseudopregnant CD-1 strain donors crossed with Vasectomized males 2 days prior^136–138^. We then collected embryos at E9.5, scored dissected embryos for percent chimerism, and selected three representative embryos for whole-embryo fate mapping.

### scRNA-seq and target site sequencing of PEtracer embryos

We prepared libraries as described by Colgan et al.^43^. Briefly, we isolated whole embryos at E9.5 and cleaned them of maternal and extraembryonic tissue. We initially screened complete embryos by fluorescence imaging to identify high-grade chimeras. All whole embryo images captured prior to scRNA-seq were captured using a Zeiss Observer 7 with a 10x air objective. We prepared images for visualization without modification using *FIJI* (v2.0.2). After this initial selection, we dissociated each embryo into single cells in TrypLE (Cat. #12604013, Invitrogen) using a pipette and carefully resuspended the cells as described above. As an additional step, we depleted red blood cells (Miltenyi Biotec Cat. #130-094-183) before filtering and concentrating cells. For each embryo, the entire cell suspension was processed for single-cell RNA sequencing (10x Genomics, Chromium GEM-X Single Cell 3’ Chip Kit v4), distributing cells across multiple reactions to remain within the maximum recommended input of cells per reaction (here, one or two reactions). We then prepared single-cell libraries following the manufacturer’s protocol, with cDNA (Step 2) and indexing amplification (Step 3) each using 10 PCR cycles. We examined primary cDNA and final libraries on a D5000 and D1000 TapeStation System, respectively (Agilent Technologies).

Following Step 2 of the 10x Genomics GEM-X workflow, we used a portion of amplified cDNA to generate target site libraries with pooled equimolar staggered forward primers and a common reverse primer, as described previously^43^. We first determined the optimal PCR1 cycle number by qPCR on a 1:8 cDNA dilution; we then performed PCR1 on undiluted cDNA, followed by a second PCR to incorporate Nextera i7 indices and sequencing adapters. We purified the final libraries by 0.9x SPRI bead cleanup and eluted them in Buffer EB. We then quantified the final libraries, standardized them to 4 nM, and pooled them for sequencing on a NovaSeq X Plus (Illumina).

#### Analyses of lineage-traced chimeric embryos

Unless otherwise noted, all lineage analyses were performed as previously described for Colgan et al.43 We analyzed reference PEtracer embryos using the same parameters and thresholds to directly compare chimeric Δ*Folr1* and reference data.

### PEtracer scRNA-seq data processing and quality control

We processed gene expression and lineage tracing cassette (LTC) libraries with *CellRanger* (v9.0.1) and used a custom *Python* (v3.11) script to call lineage marks (LMs) and integration barcodes (intBCs) for each LTC read. Quality control was performed as previously described: low-quality cells were removed by adaptive thresholding on mitochondrial and total UMI counts within each capture; LTC alleles consistent with PCR artifacts or ambient RNA were filtered on reads per UMI and support relative to the dominant allele within each cell-intBC pair; and donor, host, and doublet cells were identified using normalized LTC counts, host-specific SNPs, and allele conflict rates. We retained cells with >75% intBC detection for tree reconstruction. After transcriptional embedding (below), we additionally removed Leiden clusters enriched for allele conflicts (indicative of unresolved doublets) and cells with fewer than 10,000 total counts.

### Lineage tree reconstruction

We reconstructed lineage trees independently for each embryo using the scalable hybrid approach described previously^43^. Briefly, we converted LTC alleles into a character matrix in which rows corresponded to cells and columns to editable sites across intBCs. High-confidence splits supported by ≥3 shared edits were resolved at the top of each tree using a greedy recursive partitioning procedure, and subtrees within the resulting clades were reconstructed using the heuristic neighbor-joining algorithm implemented in *FastTree 2*^139^ (v2.2). Ancestral LM states were reconstructed using the Sankoff algorithm, edges without inferred mutations were collapsed, and branch lengths were estimated under the maximum-likelihood framework implemented in *ConvexML*^140^ (v1.0.0). Notably, branch length estimates assume consistent editing kinetics from initiation to sampling at E9.5 – an assumption that could be violated by decreased nucleotide availability in the Δ*Folr1* condition.

### Embedding and integration

Folr1-KO cells were projected into the normal E7.5-E10.0 reference *scVI*^141^ (v1.5.0) latent space using the model trained on the reference dataset^43^, applied without fine-tuning so that the latent space was defined entirely by the reference data and no Δ*Folr1*-specific adaptation was introduced. To transfer labels across datasets, we integrated reference and query latent representations using Harmony^142^ (*harmonypy*, v2.0.0), with dataset as the batch variable, and constructed a shared nearest-neighbor graph (k = 20) on the integrated representation. Harmony was used only for label transfer; all UMAP embeddings shown were computed directly from the uncorrected scVI latent space using *SCANPY*^127^ (v1.11.5), so that visualizations reflect the transcriptional state of each cell as placed in the reference space rather than a batch-corrected projection.

### Cell annotation

Rather than annotating Δ*Folr1* cells de novo, cell subtype labels were transferred from the normal reference. Subtype labels were imputed for each Δ*Folr1* chimera cell from the connectivities of its neighbors in the integrated graph, and cell type, lineage, and germ layer assignments were derived from these subtypes using the reference annotation. We imputed developmental stage using the same procedure, masking *ΔFolr1* stage labels and inferring them from neighboring wild-type cells to provide a transcriptional estimate of developmental progression independent of collection time. Labels were then refined within the Folr1-KO dataset alone: cells were re-embedded on the scVI representation and clustered with the Leiden algorithm (resolution = 2), subtypes represented by fewer than 20 cells and ambiguous parent-level annotations were removed, remaining labels were smoothed over the nearest-neighbor graph, and any cells still unlabeled were assigned by majority vote within their Leiden cluster. We grouped some PEtracer reference cell types to match annotations in the rest of the manuscript. For example, all the brain neuron types resolved in the reference are grouped into a single ‘brain neuron’ annotation here.

### Differential abundance

We compared cell type abundance to the reference at two levels of resolution. At the developmental-domain level, we calculated per-embryo compositions as the percentage of cells assigned to each domain, excluding blood. At the cell-type level, we compared per-embryo counts to the mean count across reference E9.5 replicates and expressed them as log₂ fold changes, restricting the analysis to types represented by more than 50 cells on average in the reference to exclude populations too sparse for stable ratio estimates.

### Number of extant cells over time

The number of extant cells over time was calculated as previously described^43^ by traversing each clone-level tree in half-day intervals and counting the branches present at each inferred timepoint, with each edge considered extant from the time of its parent node (inclusive) until the time of its child node (exclusive). Counts were summed across clones within each embryo and compared between Folr1-KO embryos and the mean of the wild-type E9.5 replicates.

### Identifying fate-restricted clades

We identified fate-restricted clades as previously described^43^. A node was defined as fate-restricted if at least 90% of its descendants belonged to a given fate. We selected clades using a two-pass procedure that retains the earliest qualifying ancestor for each fate while excluding its descendants from further consideration, and we handled multifurcations by assigning a parent node as the fate-restricted ancestor when multiple child branches independently qualified for the same fate. We excluded clades with a single descendant from summary statistics. For each embryo, we then computed the number of fate-restricted clades and their mean output (number of descendant cells of the restricted fate), and compared Δ*Folr1* chimeric embryos to reference E9.5 replicates.

### Ancestral linkage analysis

We calculated ancestral linkage as previously described^43^. Briefly, for each cell we identified the most recent common ancestor shared with the nearest cell of a target category and used the depth of this ancestor as a measure of relatedness, normalized by subtracting a permuted background expectation generated by randomly reassigning non-target labels while preserving the target population. We computed mean ancestral linkage independently for each embryo, at both the broad lineage and cell-type levels, and excluded categories represented by fewer than 100 cells in the reference data. Differences between genotypes were quantified by subtracting the reference matrix from the corresponding Folr1-KO matrix.

### Inference of axial position

Inferred anterior–posterior (A-P) and dorsal–ventral (D-V) positions of neural cells were assigned using the random forest models trained on spatially predictive genes fit against a published E13.5 sagittal *Stereo-seq* reference^143^ and applied here without retraining^43^. These scores should be interpreted as estimates of relative position along each axis rather than absolute anatomical location. For comparisons across the body axis, we binned cells and clades into nine intervals spanning inferred positions of 0 to 0.9, with clade positions defined as the mean inferred position of their descendant cells. Within each bin, we quantified neural cell type distributions, the number of ectoderm-restricted clades, and the mean number of cells per clade.

---

**Figure S1.**
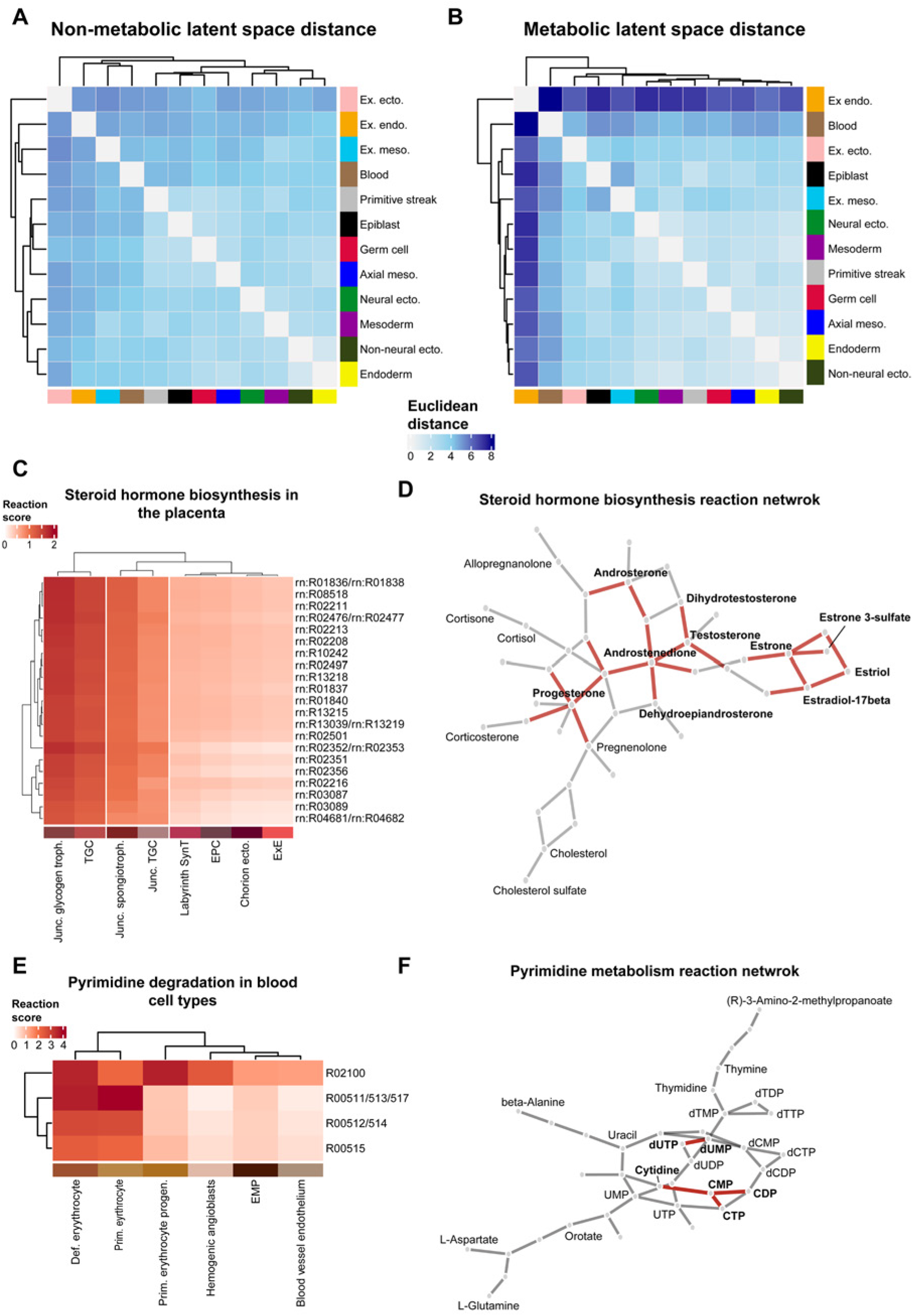
MeRN predicts metabolic reaction activity with cell-type resolution. **A,B.** Euclidean distance between lineages using average latent space positions for non-metabolic (**A**) and metabolic (**B**) predicted cell embeddings. **C**. Average extra-embryonic ectoderm reaction scores for selected reactions under the KEGG steroid hormone biosynthesis pathway, highlighting increased activity in the placental junctional zone compared to the labyrinth and chorion. Reactions were selected for variance > 0.2 within the extra-embryonic ectoderm. **D.** Network visualization of reactions in the KEGG steroid hormone biosynthesis pathway, showing the relationship between reactions (edges) and metabolites (nodes). Highlighted edges represent reactions displayed in (**C**). **E.** Average reaction scores for cell types in the blood and vascular cell lineage, highlighting activity of pyrimidine degradation reactions in differentiated erythrocytes. We curated reactions based on previously published descriptions of erythrocyte maturation^56,57^. **F.** Reaction network visualization for reactions in the KEGG pyrimidine metabolism pathway. Highlighted edges represent reactions displayed in (**E**).

**Figure S2.**
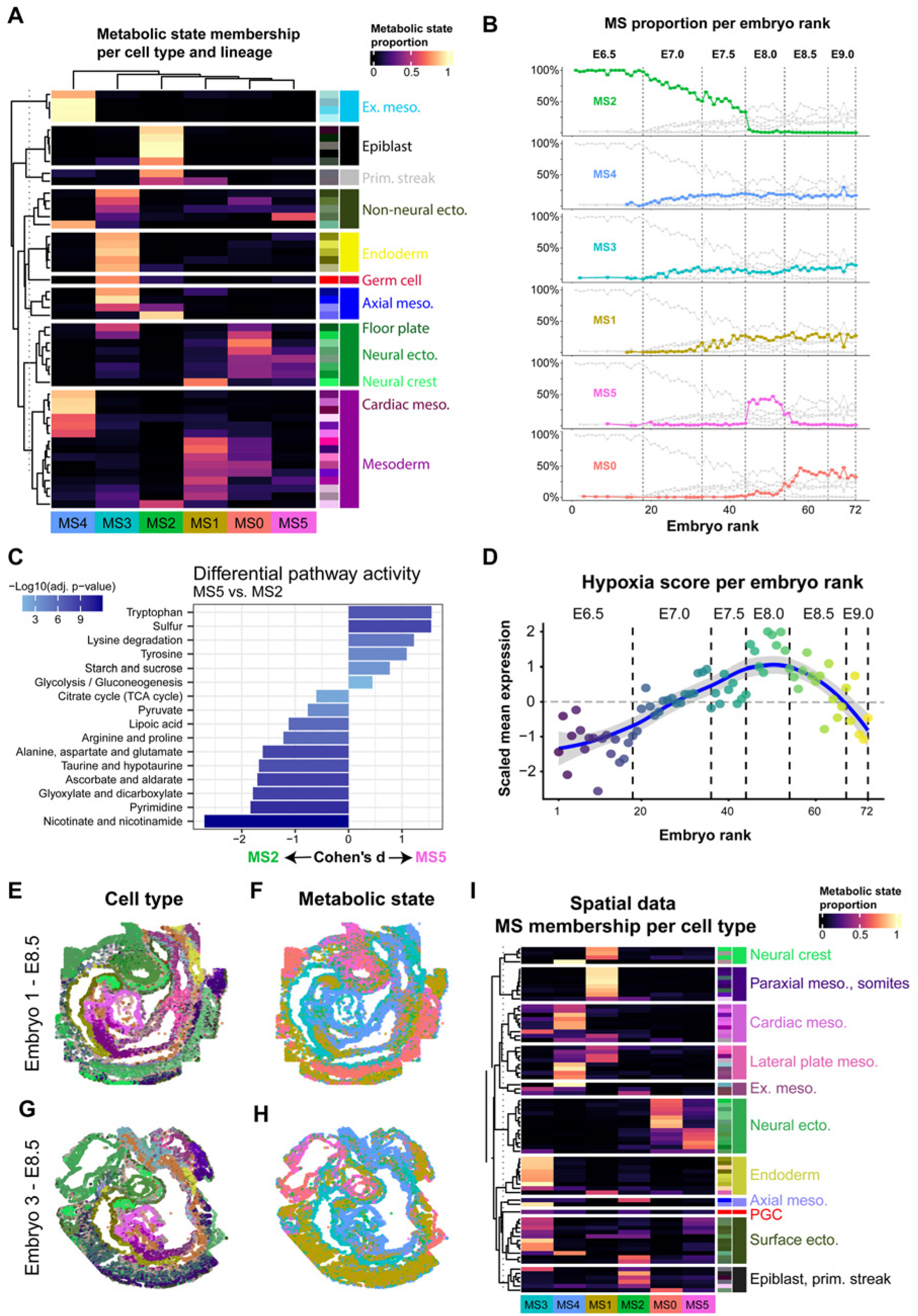
Temporal and spatial aspects of metabolic state. **A.** Heatmap of metabolic state assignments per cell type, showing the distribution of label assignments throughout the embryo. Row annotations represent individual cell types, split by overarching lineage, and columns represent metabolic states. Heat denotes the proportion of cells in a cell type assigned to each metabolic state; row values sum to 1 (100%). **B.** Proportion of metabolic state assignment per embryo, showing metabolic state abundance and dynamics over developmental time. Embryos were ranked from 1-72 according to transcriptional similarity to approximate a developmental continuum (see **Methods**). **C.** Effect size differences of preferential pathway usage between MS2 (E6.5-E7.5) and MS5 (E8.0). We curated KEGG pathways broadly related to redox homeostasis and energy metabolism that showed significantly different pathway activity (*p_adaa_* < 0.05 highlighting a preference for oxidative processes and redox protection in MS2 compared to MS5. **D.** Scaled mean expression of Hallmark hypoxia genes per embryo, displayed in ranked order as described in (**B**), highlighting maximal relative expression at E8.0, when MS5 is most abundant. **E,F,G,H.** Additional samples of seqFISH data of mouse embryos at E8.5, displayed in a sagittal section (replicates 1 and 3 from Harland et al.^75^). Cells are colored by cell type annotation (**E, G**) or predicted metabolic state (**F, H**). The spatial organization of metabolic states was consistent across all analyzed replicates. **I.** Heatmap of metabolic state assignments per cell type and lineage, as shown in (**A**), for all seqFISH data reported by Harland et al. Assignment patterns broadly matched our orthogonal data, including the association between early neural ectoderm and MS5, later neural ectoderm and MS0, cardiac mesoderm and MS4, and somitic mesoderm and MS1.

**Figure S3.**
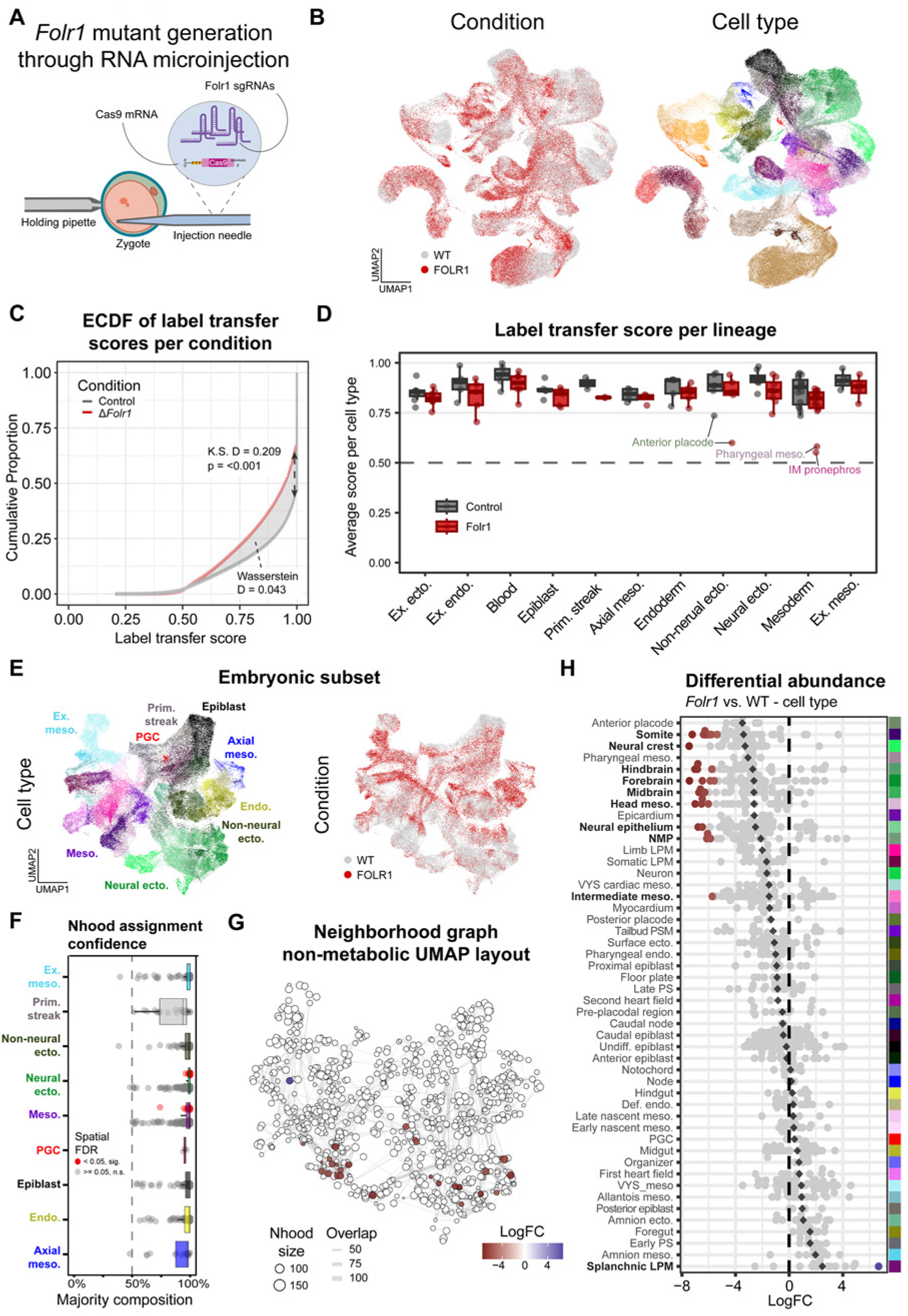
Generation and characterization of differential abundance in Δ*Folr1* embryos. **A.** Schematic of the CRISPR/Cas-9 method used to engineer zygotic crispant embryos lacking functional *Folr1*. **B.** Non-metabolic UMAP representation of the merged wild-type and Δ*Folr1* scRNA-seq data, colored by experimental condition (left) or cell type annotation (right). **C.** Empirical cumulative distribution function (ECDF) of label transfer scores, showing the difference in confidence for labels predicted for Δ*Folr1* (red) and wild-type (gray) cells, using the wild-type as a reference. Label transfer scores for each cell are cumulative across all possible cell-type labels and sum to 1.00 (100%). The gray area between curves represents the Wasserstein distance, and the double-sided arrow represents the Kolmogorov-Smirnov statistic, denoting the point of maximal vertical distance between distributions. **D.** Box plots of average label transfer score per lineage, representing confidence in label assignment. Each point represents a cell type, grouped by broad developmental lineages. Labeled points represent outliers with relatively low scores. **E.** Non-metabolic UMAP representation of the embryonic subset of merged Δ*Folr1* and wild-type cells, colored by cell type (left) and condition (right). This subset manually excludes cells in the extra-embryonic endoderm and ectoderm, as well as blood and vascular cell types. **F.** Boxplot of neighborhood composition per lineage, based on the K-nearest neighbors (KNN) graph calculated during differential abundance (DA) analysis. Points represent individual neighborhoods, positioned according to the proportion of cells belonging to the assigned label. Red points denote neighborhoods with significantly different Δ*Folr1*/wild-type (spatial FDR < 0.05). **G.** Non-metabolic UMAP layout of the KNN graph calculated using the non-metabolic latent space embeddings of wild-type and *Folr1* cells, highlighting neighborhoods with significant differential abundance and colored by log-fold-change (LogFC) calculated with Milo^87^. **H.** Differential abundance results using cell type labels, showing significant depletion of cell types in the neural ectoderm and trunk mesoderm, and enrichment of a single neighborhood in the splanchnic lateral plate mesoderm (LPM). Points represent individual neighborhoods; colored points represent significant neighborhoods, and color denotes LogFC, as in (**G**).

**Figure S4.**
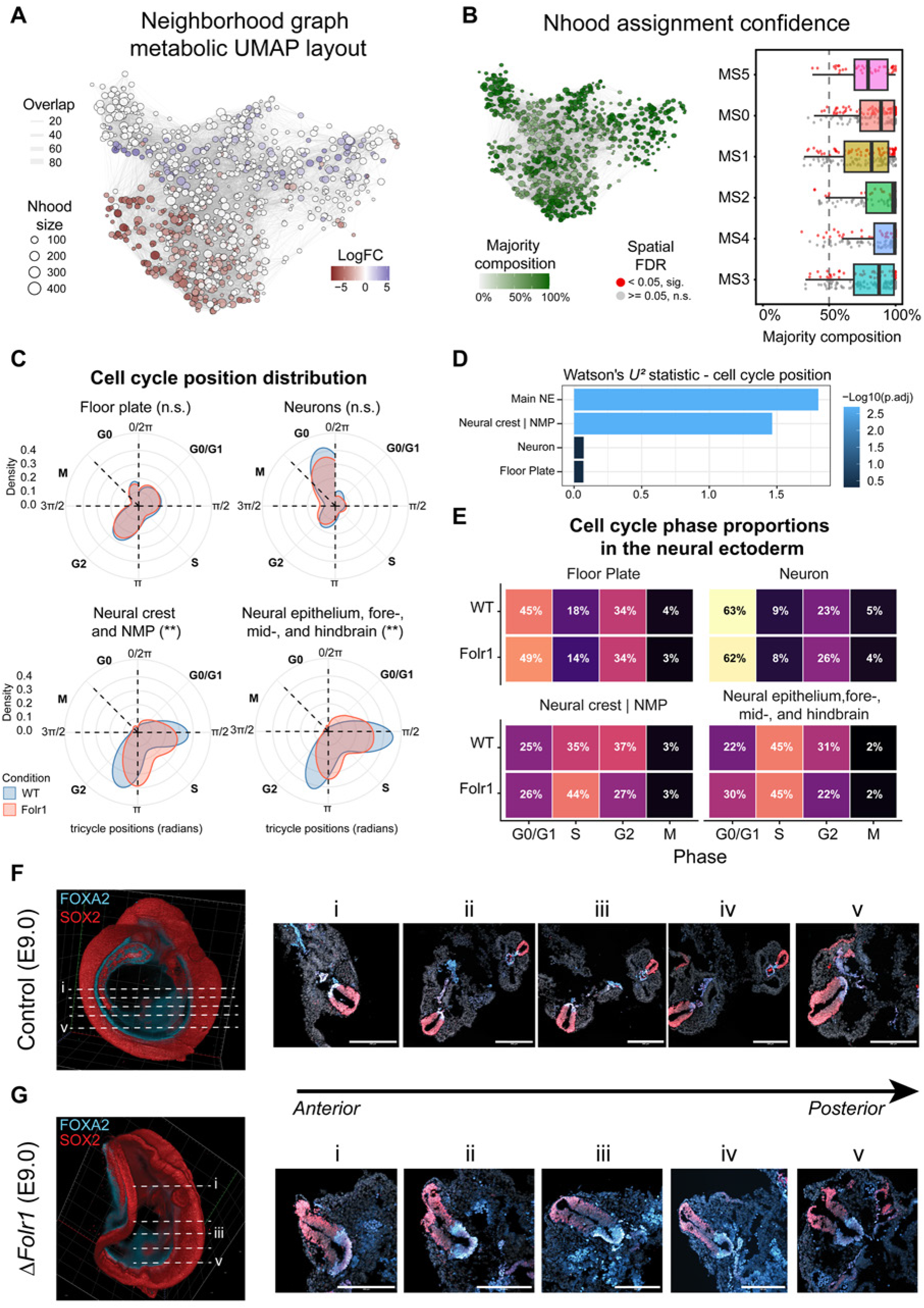
Cell depletion dynamics correlate with metabolic state and cell cycle dynamics. **A,B.** Metabolic UMAP layout of the KNN graph calculated using the metabolic latent space embeddings of wild-type and *Folr1* cells, highlighting neighborhoods with significant differential abundance and colored by LogFC calculated with Milo^87^ (**A**) or proportion of cells in the assigned (majority) label (**B**). The box plot shows the proportion of cells in each neighborhood belonging to the assigned label, split by metabolic state. Colored points denote significant neighborhoods (spatial FDR < 0.05). **C,D.** Polar coordinate representation of neural ectoderm cell cycle position distributions calculated with tricycle^94^, divided into four classes and displayed in (**C**). Significance was calculated using 999 permutations of Watson’s two-sample test of homogeneity and adjusted with the Benjamini-Hochberg correction (**: *p_adaa_* < 0.01). Panel (**D**) also shows the adjusted p-value and Watson’s U2 statistic per group. Main NE refers to the group containing neural epithelium, fore-, mid-, and hindbrain. **E.** Heatmap of cell cycle phase proportions per group, assigned according to cell cycle position, and highlighting the accumulation of neural crest and NMP cells in S phase, and main neural ectoderm cells in G0/G1 in response to genetic ablation of *Folr1*. **F,G.** Representative images of light-sheet microscopy sections of the neural tube, showing transverse sections of the neural tube and highlighting the expansion of the floor plate domain in Δ*Folr1* (**G**) compared to wild-type (**F**). Immunostaining shows markers for neural ectoderm (SOX2, red) and floor plate and axial mesoderm (FOXA2, blue). Dashed lines show the approximate location of the transverse sections, numbered in ascending order along the anterior-posterior axis. Scale bars: 200 µm.

**Figure S5.**
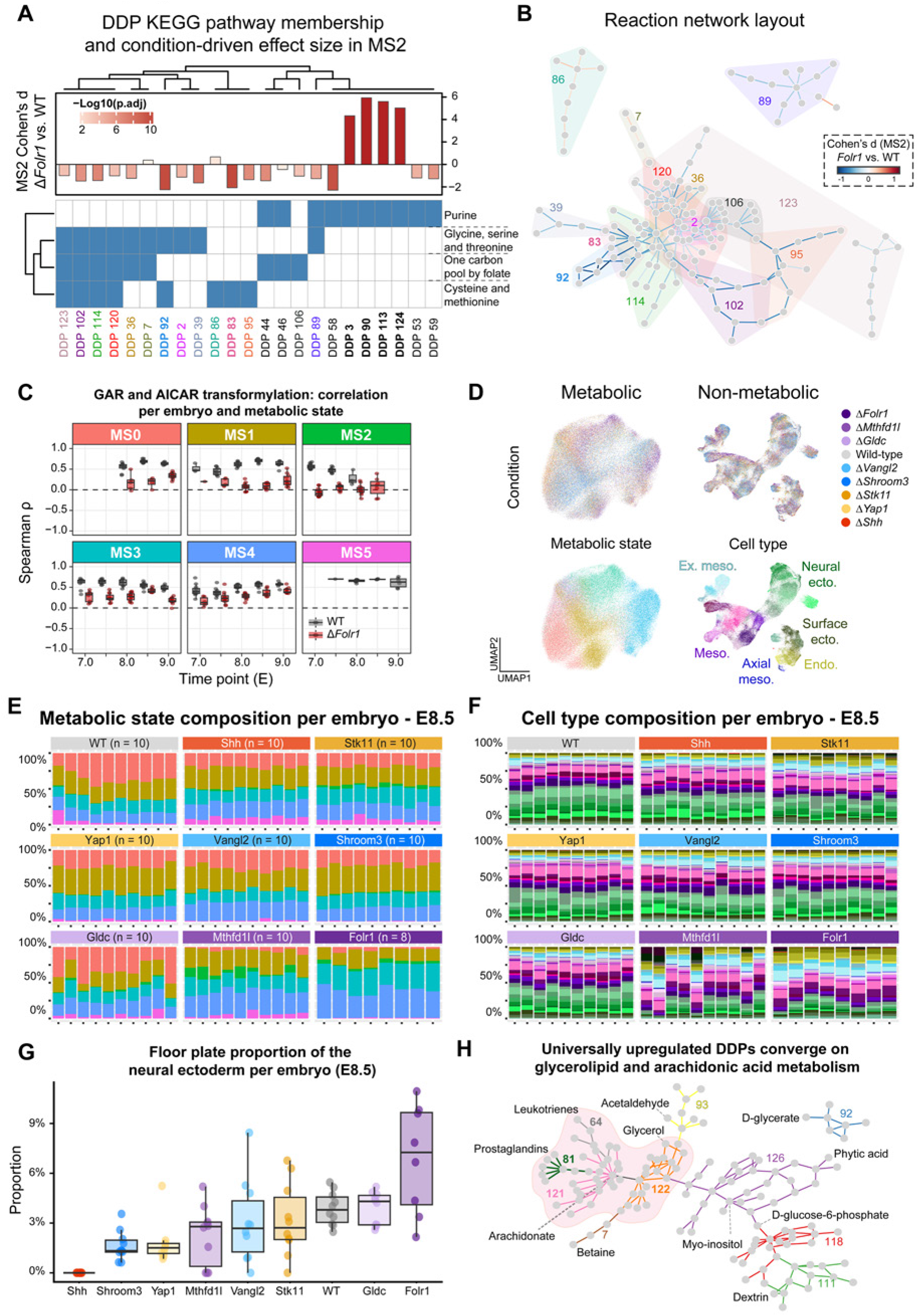
MeRN predicts metabolic determinants of neural tube defect etiology. **A.** Membership heatmap of DDPs in select folate-related KEGG pathways, highlighting the outsized effect of *Folr1* disruption on purine metabolism. **B.** Reaction network layout showing non-purine metabolism reactions displayed in (**A**), colored by reaction Cohen’s d between WT and Folr1 in MS2. **C.** Box plots of Spearman correlation between GAR and AICAR transformylation reaction scores per embryo and metabolic state over time, indicating a global loss in correlation between folate-dependent reactions required for *de novo* purine biosynthesis. Each point represents an embryo; box plots are colored by condition. **D.** UMAP representations of the merged scRNA-seq data for 78 mutant embryos and 10 wild-type embryos at E8.5. UMAPs were generated using metabolic (left) or non-metabolic (right) latent space embeddings, and colored by experimental condition (top), metabolic state (bottom left), and cell type (bottom right). **E,F.** Bar plots of embryo composition. Each bar represents one embryo; slices are colored according to metabolic state (**E**) and cell type (**F**) labels, showing broad condition-driven compositional dynamics. **G.** Box plots denoting the proportion of the neural ectoderm represented by the floor plate, colored by experimental condition, and highlighting both the absence of the floor plate in Δ*Shh* crispant embryos and the enrichment of this ventral organizer in Δ*Folr1*. Points denote individual embryos. **H.** Reaction network layout showing the common group of DDPs upregulated across all mutant conditions (**64, 81, 121, 122**, highlighted). Notably, these DDPs are connected within glycerolipid and arachidonic acid metabolism, suggesting a common metabolic response to genetic insults associated with neural tube defects. The displayed DDPs include reactions within the glycerolipid, glycerophospholipid, or arachidonic acid metabolism KEGG pathways (**rn00561**, **rn00564**, and **rn00590**, respectively).

**Figure S6.**
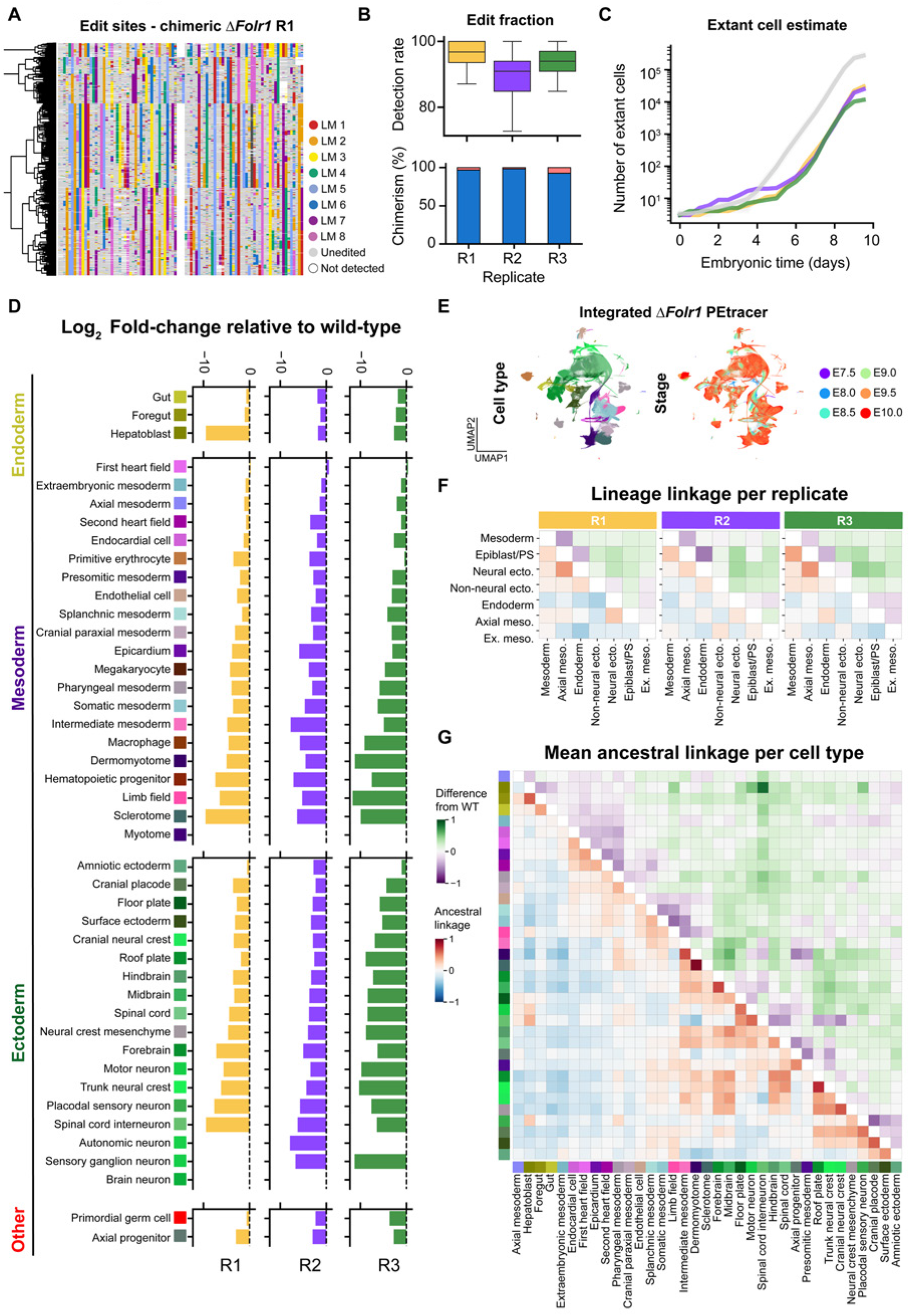
Chimeric Δ*Folr1* embryos display disrupted composition. **A.** Representative reconstructed lineage tree and character matrix for chimeric Δ*Folr1* replicate 1 (R1). Character matrix columns represent editable sites (3 per barcoded lineage-tracing cassette, 105 total sites) and are colored by each cell’s edit state. LM = lineage marker. **B.** Box plots quantifying the per-cell detection rate for lineage-tracing cassettes (top) and bar plots denoting the proportion of the embryo represented by donor (blue) and host (pink) cells for each chimeric mutant replicate (bottom). R = Replicate. **C.** Estimated number of extant cells as a function of inferred developmental time based on branch length estimates, colored by replicate as in (**B**). Gray line represents the E9.5 reference mean. **D.** Δ*Folr1* log_2_ fold-change per cell type relative to the E9.5 reference, calculated per replicate chimeric mutant embryo. Missing bars denote complete absence in the Δ*Folr1* condition. **E.** UMAP representation of the integrated Δ*Folr1* and E7.5-E10.0 reference PEtracer data, colored by cell type as displayed in (**D**) (left) and by developmental stage. **F,G.** Mean pairwise ancestral linkage (**Methods**) at the level of broad lineages (**F**) and cell type (**G**). Below the diagonal (red-blue) indicates Δ*Folr1* linkage, with the mean across replicates in **G**, and above the diagonal (green-purple) indicates the relative difference from the mean E9.5 reference linkage between the same categories.

